# TGF-β–TIF1γ signaling restrains CD8⁺ T cell expansion while preserving cytotoxic competence

**DOI:** 10.64898/2026.11.05.764687

**Authors:** Jesús Daniel Zambrano-Romero, Eugenio Contreras-Castillo, Verónica Yutsil García-Rasilla, Cleofas Marcial-Medina, Diana Berenice Ríos-Ramírez, José Luis Ramos-Balderas, Javier Orozco-Córdoba, Luis Tenorio-Hernández, Alfredo Ibarra-Sánchez, Diego Delgado-Zaldívar, Harumichi Ishigame, Maria Carolina Amezcua Vesely, Claudia González-Espinosa, Florencia Rosetti, José C. Crispín, Mayra Furlan-Magaril, Vesa Kaartinen, Richard Flavell, Paula Licona-Limón

**Affiliations:** Departamento de Biología Celular y Desarrollo, Instituto de Fisiología Celular, Universidad Nacional Autónoma de México, 04510 Mexico City, Mexico; Departamento de Farmacobiología y Centro de Investigación sobre el Envejecimiento, Centro de Investigación y de Estudios Avanzados, Unidad Sede Sur, Mexico City 14330, Mexico; Departamento de Genética Molecular, Instituto de Fisiología Celular, Universidad Nacional Autónoma de México, 04510 Mexico City, Mexico; Departamento de Inmunología y Reumatología, Instituto Nacional de Ciencias Médicas y Nutrición Salvador Zubirán. 14080, Mexico City, Mexico; Escuela de Medicina y Ciencias de la Salud, Tecnológico de Monterrey, 64700 Monterrey, Mexico; Laboratory for Tissue Dynamics, RIKEN Center for Integrative Medical Sciences, Yokohama, Kanagawa 230-0045, Japan; Near-Infrared Photo-Immunotherapy Research Institute, Kansai Medical University, Hirakata, Osaka 573-1010, Japan; Centro de Investigación en Bioquímica Clínica e Inmunología, CIBICI-CONICET, Facultad de Ciencias Químicas, Universidad Nacional de Córdoba, 5000, Argentina; Department of Biologic and Materials Sciences, University of Michigan School of Dentistry, Ann Arbor, MI, USA; Yale University, New Haven, CT, USA

## Abstract

Transforming growth factor-β (TGF-β) exerts complex and often opposing effects on CD8^+^ T cell biology, yet the molecular mechanisms that distinguish these functions remain poorly understood. Here, we identify TIF1γ (TRIM33) as a critical signal threshold regulator that couples TGF-β signaling to CD8^+^ T cell activation and cytotoxic function. In the presence of TGF-β, TIF1γ-deficient CD8^+^ T cells bypass canonical suppressive signals and acquire a transcriptional program characterized by enhanced activation and proliferation. Despite this hyperactivated state, TIF1γ-deficient cells fail to acquire full effector function, exhibiting markedly reduced Granzyme B and Perforin expression under TGF-β signaling. We confirmed this functional uncoupling *in vivo* using a secondary *Listeria monocytogenes* challenge model, in which TIF1γ-deficient CD8^+^ T cells underwent robust expansion yet failed to efficiently control bacterial infection because of impaired cytotoxic activity. In a lymphopenic homeostatic proliferation model, TIF1γ deficiency disrupted cellular quiescence, leading to excessive proliferation and spontaneous effector differentiation. Consistent with these findings, loss of TIF1γ accelerated B16 melanoma growth and was associated with reduced Perforin expression in tumor-infiltrating CD8^+^ T cells. Together, these findings identify TIF1γ as a key regulator of TGF-β–mediated CD8^+^ T cell responses, uncoupling activation from cytotoxic differentiation and revealing a mechanism by which TGF-β independently regulates proliferative and effector programs.

## Introduction

CD8⁺ T cells are central mediators of protective immunity against intracellular pathogens and cancer. Upon antigen encounter, naïve CD8⁺ T cells (T_N_) undergo extensive clonal expansion and differentiate into cytotoxic effector (T_E_) and long-lived memory (T_M_) subsets ^1,2^. This process is tightly regulated by microenvironmental cues, among which transforming growth factor-β (TGF-β) functions as a context-dependent regulator of CD8⁺ T cell fate. TGF-β restrains the activation and proliferation of naïve CD8⁺ T cells, limits the cytotoxic function of effector cells to prevent immunopathology, and promotes the differentiation of tissue-resident memory (T_RM_) and exhausted (T_PEX_/T_EX_) populations ^3–7^.

Canonical TGF-β signaling is mediated by TGFβRI/TGFβRII receptor complexes that phosphorylate Smad2 and Smad3, which subsequently associate with Smad4 to regulate gene transcription ^8^. In CD8⁺ T cells, SMAD4 has been implicated in the regulation of genes encoding negative regulators of TGF-β signaling, including *Ski*, *Skil*, *Smad7*, and *Smurf2*, as well as genes involved in tissue residency and migration, such as *Itgae*, *Klrg1*, *Sell*, *Cxcr3*, and *Eomes*, and effector function, including *Tbx21* and *Gzma* ^9–11^. However, the effects of SMAD4-dependent regulation of these genes can vary depending on the extent to which SMAD4 is required for TGF-β responses and on the activation and differentiation state of CD8⁺ T cells, complicating the interpretation of SMAD4-dependent functions in *in vivo* mouse models ^11–14^. This complexity raises the possibility that additional TGF-β-regulated pathways contribute to CD8⁺ T-cell responses independently of SMAD4.

An alternative, SMAD4-independent branch of TGF-β signaling involves Transcriptional Intermediary Factor 1γ (TIF1γ; also known as TRIM33), which can associate with activated SMAD2/3 to regulate gene expression ^15,16^ and could lead to a different gene transcriptional profile than that driven by Smad4. TIF1γ has emerged as an important regulator of immune cell differentiation and function across multiple lineages. During T-cell development, TIF1γ acts downstream of TGF-β to promote invariant natural killer T (iNKT) cell maturation by supporting the survival of early thymic precursors ^17^. In peripheral CD4⁺ T cells, TIF1γ promotes TH17 differentiation by regulating the balance between IL-10 and IL-17A expression ^18^, and we recently demonstrated that it preserves regulatory T (Treg) cell stability during inflammation by promoting β-catenin degradation and repressing the acquisition of pro-inflammatory programs ^19^. Beyond T cells, TIF1γ also limits inflammatory responses by repressing *Ifnb1* expression in macrophages following LPS stimulation ^20^, functions as an E3 ubiquitin ligase that targets HIV-1 integrase in infected CD4⁺ T cells ^21^, and supports the survival of plasmacytoid and conventional dendritic cells through regulation of *Irf8* and *Bcl2l11* expression ^22,23^.

Despite the broad influence of TGF-β on CD8^+^ T cell biology, how this pleiotropic cytokine dictates distinct and often opposing transcriptional outcomes remains a fundamental question. Here, we demonstrate that TIF1γ functions as a critical molecular switch within TGF-β signaling, segregating cellular proliferation and activation from cytotoxic effector capacity to shape functional CD8^+^ T cell fate. Transcriptomic analyses of wild-type and TIF1γ-deficient CD8⁺ T cells revealed that TIF1γ selectively mediates the TGF-β-dependent repression of activation and cell-cycle programs while preserving cytotoxic differentiation. As a consequence, TIF1γ-deficient CD8⁺ T cells exhibited enhanced TCR sensitivity and exaggerated homeostatic proliferation *in vivo*. Surprisingly, this hyperproliferative state did not improve immune protection. Instead, loss of TIF1γ impaired control of *Listeria monocytogenes* infection and accelerated B16 melanoma progression, demonstrating that increased expansion can be uncoupled from effective cytotoxic function. Together, our findings identify TIF1γ as a key molecular determinant of TGF-β signaling, coordinating CD8⁺ T cell expansion with protective effector function.

## Results

### TIF1γ is required for TGF-β-mediated control of CD8⁺ T cell proliferation and differentiation

To explore the role of TIF1γ in T cell immunity, we first assessed its expression profile across peripheral lymphocyte subsets isolated from wild-type (WT) mice. Quantitative PCR analysis revealed that *Tif1g* was expressed at comparable levels in CD4⁺ T cells, regulatory T cells (Tregs), and CD8⁺ T cells under steady-state conditions (Supplementary Fig. 1A). TIF1γ operates downstream of TGF-β in other immune lineages, yet regulation of the *Tif1g* gene transcription by TGF-β has not been reported in CD8^+^ T cells. To assess this, we isolated naïve CD8⁺ T cells from *Tif1g^fl/fl^*(WT, hereinafter) mice and stimulated them with anti-CD3 and anti-CD28 in the presence or absence of exogenous TGF-β. Notably, TGF-β exposure led to a significant upregulation of *Tif1g* expression in activated CD8⁺ T cells after five days in culture (Supplementary Fig. 1B), establishing *Tif1g* as a TGF-β-inducible gene during CD8⁺ T cell activation.

To functionally dissect the cell-intrinsic role of TIF1γ in mature CD8⁺ T cells, we crossed *Tif1g^fl/fl^*mice with *Cd8a-Cre* mice to generate conditional knockout animals (*Cd8a-Cre Tif1g^fl/fl^*, hereafter referred to as TIF1γ KO). Immunophenotypic analyses of adult mice at steady state revealed that conditional deletion of TIF1γ did not alter the frequencies of peripheral CD8⁺ T cells, nor did it disturb the distribution of naïve (T_N_; CD62L^+^ CD44^-^), memory (T_M_; CD62L^+^ CD44^+^), effector (T_EFF_; CD62L^-^ CD44^+^), and transitory (T_TRAN_; CD62L^-^ CD44^-^) CD8^+^ T cells within the spleen or peripheral, salivary, and mesenteric lymph nodes (Supplementary Fig. 1C–G). Thus, TIF1γ deletion does not affect peripheral CD8⁺ T-cell homeostasis.

To define the contribution of TIF1γ to TGF-β-mediated regulation of CD8⁺ T cells, naïve WT and TIF1γ-deficient cells were activated in the presence or absence of TGF-β and IL-2. We then assessed proliferation, survival, and differentiation. Functional assessment revealed that TIF1γ KO cells exhibited a marked resistance to the anti-proliferative effects of TGF-β after four days of culture in the absence of IL-2 (Fig. 1A). Remarkably, the addition of exogenous IL-2 completely rescued this phenotype, aligning the proliferative capacity of WT and TIF1γ KO cells (Fig. 1B), demonstrating that IL-2 signaling is sufficient to recover the TIF1γ-dependent inhibition of CD8^+^ T cell proliferation. Given that TIF1γ can restrict cell proliferation by promoting the degradation of β-catenin in tumor cells ^24^ and during inflammatory Treg instability ^19^, we investigated whether this axis was shared in CD8⁺ T cells. Kinetic analysis of intracellular β-catenin accumulation showed that TGF-β significantly reduced the frequency of CTV^-^ β-catenin^+^ cells across days in both phenotypes in the absence of IL-2 (Supplementary Fig. 1H). Additionally, β-catenin MFI within the positive population remained unchanged across genotypes and treatments (Supplementary Fig. 1I). This suggests that β-catenin is not involved in the proliferative phenotype observed in TIF1γ-deficient CD8^+^ T cells.

**Fig. 1.**
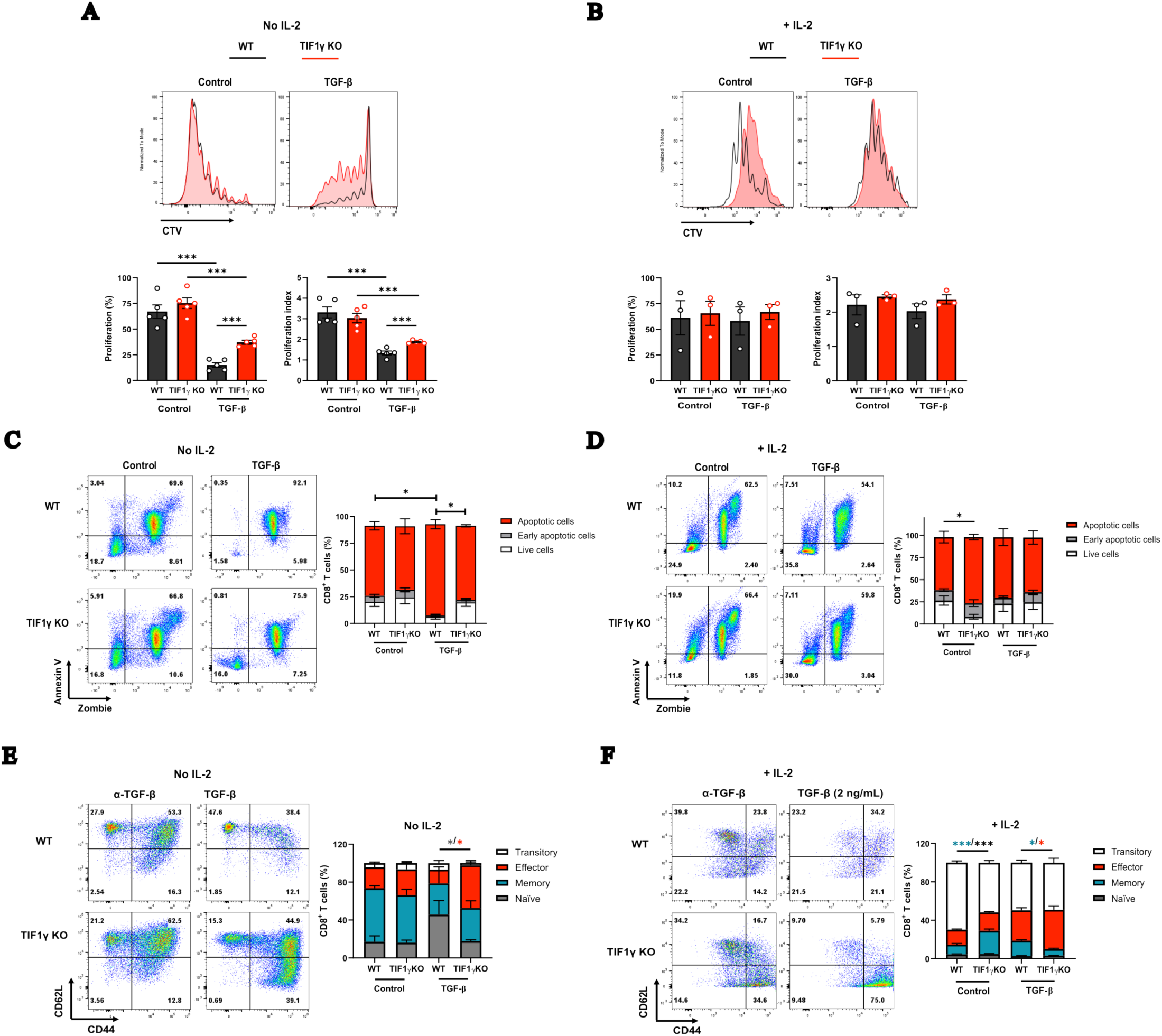
TIF1γ deficiency bypasses TGF-β-mediated antiproliferative, pro-apoptotic, and differentiation blocks in activated CD8^+^ T cells. Naïve WT and TIF1γ KO CD8^+^ T cells were isolated and activated *in vitro* using plate-bound anti-CD3 (2 µg/mL) and anti-CD28 (1 µg/mL) for 4 days in the presence of neutralizing anti-TGF-β (2 µg/mL) or recombinant mouse TGF-β1 (2 ng/mL). (A, B) Representative flow cytometry histograms of CellTrace Violet (CTV) dilution (top), along with cumulative frequencies of proliferating cells and calculated proliferation indices (bottom) maintained in the absence (A) or presence (B) of exogenous IL-2 (20 ng/mL). (C, D) Representative pseudocolor dot plots and cumulative percentages of late apoptotic (Annexin V^+^ Zombie^+^), early apoptotic (Annexin V^+^ Zombie^−^), and viable (Annexin V^−^ Zombie^−^) cells cultured without (**C**) or with (**D**) exogenous IL-2. (**E, F**) Representative flow cytometry plots and cumulative frequencies of differentiation subsets, classified as naïve (CD62L^+^CD44^−^), central memory (CD62L^+^CD44^+^), effector memory (CD62L^−^CD44^+^), and transitional (CD62L^−^CD44^−^) CD8^+^ T cells, in the absence (**E**) or presence (**F**) of exogenous IL-2. Asterisk colors correspond to statistically significant differences within matching cell populations or fractions (bar colors). Data represent the mean ± SEM of three or five independent experiments. Statistical significance was determined by a two-tailed unpaired Student’s t-test or two-way ANOVA (with post-hoc testing where applicable). \**P < 0.05; ***P < 0.001*.

We next examined the impact of TIF1γ on TGF-β-induced apoptosis. Although both WT and TIF1γ KO cells displayed baseline activation-induced cell death, exogenous TGF-β significantly accelerated apoptosis in WT cells in the absence of IL-2; this increase was absent in TIF1γ-deficient cells (Fig. 1C). Similar to our proliferation findings, the addition of IL-2 neutralized the pro-apoptotic effects of TGF-β in WT cells, thereby eliminating any genotypic differences (Fig. 1D). Beyond proliferation restraint and apoptosis induction, TGF-β signaling is critical for restraining effector differentiation. Consistently, TGF-β limited the emergence of effector-phenotype (CD62L^-^ CD44^+^) cells in activated WT cultures; however, TIF1γ KO cells resisted this restriction, driving robust effector differentiation regardless of IL-2 availability (Fig. 1E, F). In contrast, in the absence of TGF-β but with IL-2, TIF1γ KO cells showed elevated baseline apoptosis alongside an enriched proportion of memory-phenotype CD8^+^ T cells (CD62L^+^ CD44^+^) (Fig. 1D, F). Together, these data demonstrate that TIF1γ is required for TGF-β-mediated regulation of CD8⁺ T cell proliferation, survival, and differentiation.

To assess whether these defects stemmed from a loss of TGF-β sensitivity, we examined the expression dynamics of the type I TGF-β receptor (TGFβRI). Naïve WT and TIF1γ KO cells displayed similar basal surface levels of TGFβRI (Supplementary Fig. 1J). Notably, during in vitro activation without IL-2, TGF-β exposure sustained surface TGFβRI expression in WT but not in TIF1γ KO cells at day 5 in culture (Supplementary Fig. 1K, L). Although this significant difference in TGFβRI surface availability occurred after the onset of the proliferative, survival, and differentiation phenotypes, a gradual loss of TGFβRI surface expression was observed over days in culture, suggesting that TIF1γ is involved in maintaining TGFβRI on the cell membrane during long-term TGF-β stimulation. Together, these findings establish TIF1γ as a critical molecular mediator of the antiproliferative and inhibitory effects of TGF-β on effector differentiation in CD8⁺ T cells. However, whether the enhanced expansion and effector differentiation observed in the absence of TIF1γ translate into improved cytotoxic function remained unknown.

### TIF1γ preserves CD8⁺ T cell cytotoxic function under TGF-β signaling

We next asked whether TIF1γ contributes to the suppression of CD8⁺ T-cell cytotoxicity by TGF-β. To delineate this, we immunized polyclonal WT and TIF1γ KO mice with ovalbumin, followed by the adoptive transfer of target splenocytes pulsed with the SIINFEKL peptide 3 weeks after immunization (Fig. 2A). Interestingly, endogenous antigen-specific CD8⁺ T cells in TIF1γ KO mice displayed a significantly reduced capacity for specific target lysis *in vivo* (Fig. 2B). Further examination of endogenous CD8^+^ T cells showed that the frequencies of GzmB^+^, but not Prf1^+^, TIF1γ KO cells were lower than their WT counterparts (Fig. 2C, D). Moreover, TIF1γ KO cells exhibited lower levels of Prf1 and GzmB compared to WT cells (Fig.2E, F), demonstrating impaired cytotoxic function in TIF1γ-deficient CD8⁺ T cells.

**Fig. 2.**
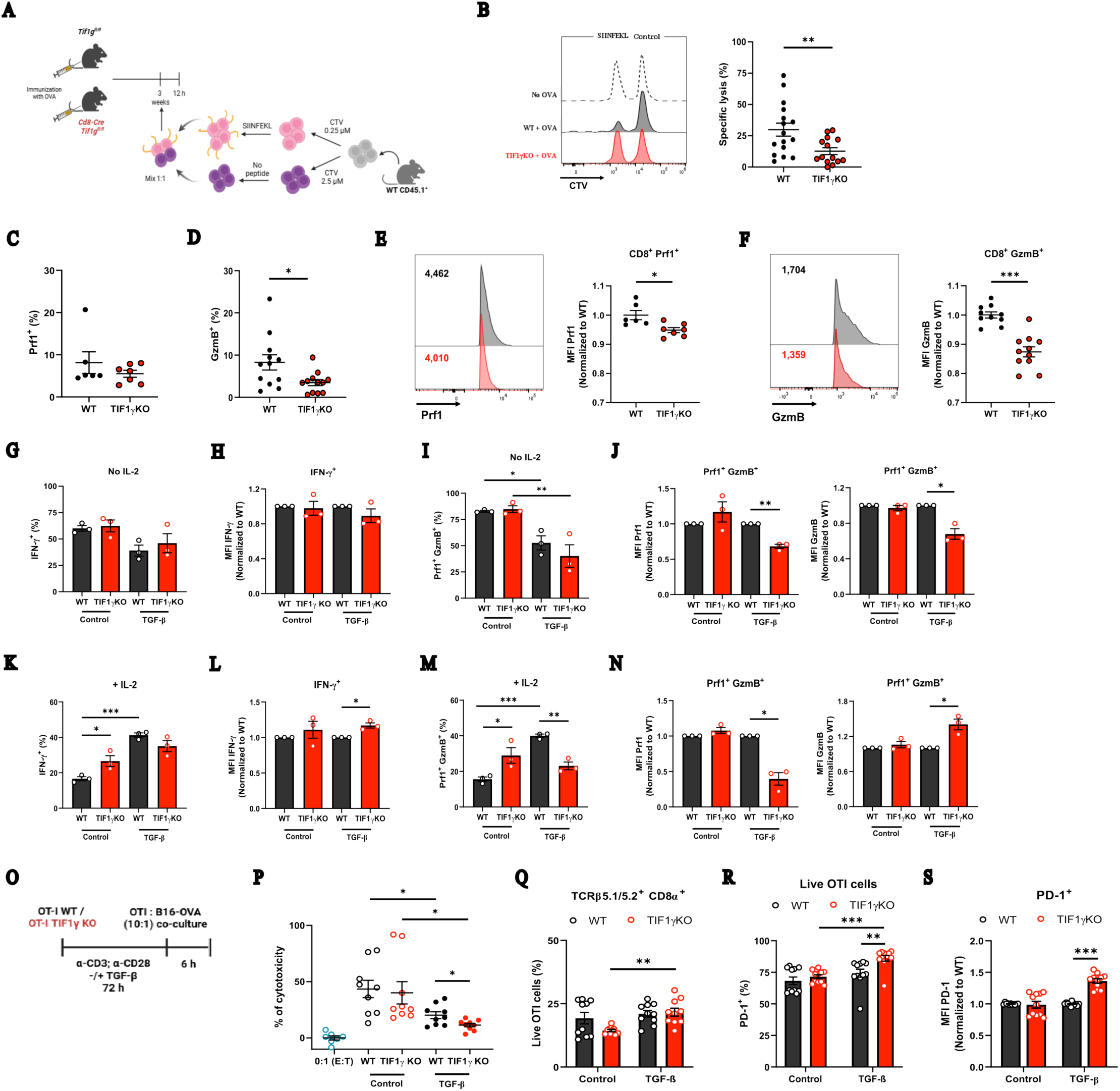
TIF1γ-deficient CD8^+^ T cells exhibit exacerbated inhibition of the cytotoxic function by TGF-β. (**A**) Schematic overview of the *in vivo* cytotoxicity assay. (**B)** Representative flow cytometry histograms and calculated frequencies of specific lysis of target splenocytes pulsed with SIINFEKL peptide and adoptively transferred into recipient mice. (**C**) Frequencies of Prf1^+^ and (**D**) GzmB^+^ CD8^+^ T cells. (**E, F**) Representative histograms and calculated frequencies of Prf1^+^ and GzmB^+^ CD8^+^ T cells. (**G–J**) Naïve WT and TIF1γ KO CD8⁺ T cells were isolated and activated *in vitro* with plate-bound anti-CD3 (2 µg/mL) and anti-CD28 (1 µg/mL) for 4 days in the presence of neutralizing anti-TGF-β (2 µg/mL) or TGF-β1 (2 ng/mL). Panels (**G–J**) display cultures maintained in the absence of exogenous IL-2: frequencies of IFN-γ^+^ CD8⁺ T cells (**G**), normalized MFI of IFN-γ (**H**), frequencies of Prf1^+^ GzmB^+^ cytotoxic CD8⁺ T cells (**I**), and normalized MFI of Prf1 and GzmB (**J**). Panels (**K–N**) display matching parameters for cultures supplemented with recombinant human IL-2 (20 ng/mL): frequencies of IFN-γ^+^ CD8⁺ T cells (**K**), normalized MFI of IFN-γ (**L**), frequencies of Prf1^+^ GzmB^+^ CD8⁺ T cells (**M**), and normalized MFI of Prf1 and GzmB (**N**). (**O**) Schematic experimental design of the *in vitro* killing assay utilizing control or TGF-β1-pretreated WT and TIF1γ KO OT-I cytotoxic T lymphocytes (CTLs) co-cultured with B16-OVA melanoma target cells at a 10:1 ratio. (**P**) Specific cytotoxicity percentage determined by extracellular LDH release. (**Q**) Frequency of viable remaining OT-I CTLs (TCR*β*5.1/5.2^+^ CD8*α*^+^ Annexin V^−^ Zombie^−^). (**R, S**) Frequency (**R**) and normalized MFI (**S**) of PD-1 surface expression on viable OT-I CTLs. All MFI values are presented as normalized values to their respective WT counterparts within each experiment. Data represent the mean ± SEM of two (**C, E**), three (**D, F, G–S**) or four (**B**) independent experiments. For panel B: WT, n = 16; TIF1γ KO, n = 14. For panels C and E: WT, n = 6; TIF1γ KO, n = 7. For panels D and F: WT, n = 12; TIF1γ KO, n = 12. For panels P–S: WT, n = 9; TIF1γ KO, n = 9. Statistical significance was determined by a two-tailed unpaired Student’s t-test or a Mann-Whitney U test (depending on data normality) for frequencies and absolute values, or a one-sample t-test for normalized MFI values. \**P < 0.05; \**\**P < 0.01; ***P < 0.001*.

We then systematically profiled the production of the core effector molecules IFN-γ, GzmB, and Prf1 in activated WT and TIF1γ KO cells stimulated in the presence or absence of TGF-β, in cultures without exogenous IL-2. Under these baseline polarizing conditions, the frequency and expression levels of IFN-γ remained equivalent between genotypes (Fig. 2G, H). However, while TGF-β expectedly restricted the proportion of Prf1^+^ GzmB^+^ cytotoxic cells in both groups (Fig. 2I), TIF1γ KO cells exhibited a severe reduction in per-cell levels (measured by MFI) of both Prf1 and GzmB compared to WT controls (Fig. 2J). These findings indicate that TIF1γ is required to maintain Prf1 and GzmB expression during TGF-β signaling.

Because IL-2 opposes TGF-β-mediated suppression of effector molecule expression ^3,25^, we next asked whether adding exogenous IL-2 to cultures could rescue the exacerbated TGF-β-mediated repression of cytotoxic molecule expression observed in TIF1γ KO cells. In the absence of TGF-β, TIF1γ KO cell cultures exhibited elevated baseline frequencies of IFN-γ^+^ and Prf1^+^ GzmB^+^ populations compared with WT cell cultures (Fig. 2K, M), indicating that TIF1γ deficiency alters the response to IL-2. However, under these conditions, the levels of IFN-γ, Prf1, and GzmB did not differ between genotypes (Fig. 2L, N). Under combined TGF-β and IL-2 stimulation, although the frequency of IFN-γ^+^ cells was similar between WT and TIF1γ KO cells, the latter exhibited higher levels of IFN-γ (Fig. 2K, L). Notably, the combination of TGF-β and IL-2 induced a higher frequency of the Prf1^+^ GzmB^+^ population in WT cells, but not in TIF1γ KO cells, compared to control conditions (Fig. 2M). This translated into a lower frequency of cytotoxic T cells in TIF1γ KO cell cultures under TGF-β and IL-2 stimulation (Fig. 2M). As with IFN-γ, GzmB expression was largely restored or even elevated in TIF1γ KO cells relative to WT counterparts (Fig. 2L, N), suggesting that IL-2 signaling bypasses TIF1γ to counteract TGF-β-mediated repression of these loci. Conversely, Prf1 expression remained severely blunted in TIF1γ KO cells even in the presence of IL-2 (Fig. 2N).

To determine whether the TIF1γ deletion directly impairs immediate cytolytic efficiency against malignancies, we performed an *in vitro* killing assay by co-culturing equal numbers of OT-I WT or OT-I TIF1γ KO cells with OVA-expressing B16 (B16-OVA) melanoma cells in the absence of IL-2 (Fig. 2O). TGF-β-pretreated OT-I TIF1γ KO cells exhibited a severe defect in their cytolytic capacity compared to OT-I WT cells (Fig. 2P). This functional failure was not driven by unequal numbers of live OT-I cells between genotypes at the end of the culture (Fig. 2Q). Instead, we observed a higher frequency of PD-1^+^ cells and higher levels of PD-1 expression in TIF1γ KO co-cultures (Fig. 2R, S), indicating that TIF1γ deficiency drives a dysfunctional state that compromises the anti-tumor cytotoxic capacity of CD8⁺ T cells. These findings reveal a striking disconnect between activation and cytotoxic function in TIF1γ-deficient CD8⁺ T cells, suggesting that TIF1γ selectively regulates distinct transcriptional programs downstream of TGF-β.

### TIF1γ defines a transcriptional branch of TGF-β signaling controlling CD8⁺ T cell activation

While the global transcriptional targets of TGF-β in CD8⁺ T cells are well documented ^3,4^, the specific gene networks regulated through the alternative TIF1γ branch remain completely undefined. To define the transcriptional programs regulated by TIF1γ downstream of TGF-β, we performed RNA-seq on activated WT and TIF1γ-deficient CD8⁺ T cells cultured in the presence or absence of TGF-β. Cultures were maintained without IL-2 to isolate TGF-β-dependent effects. In WT cells, TGF-β exposure induced a robust transcriptional overhaul, significantly upregulating 4,315 genes, including canonical targets such as *Itgae*, *Tgfbr1*, *Ccr8*, *Ccr6*, *Ccl20*, and *Cd276*, while downregulating 4,702 genes associated with T cell activation, effector functions, and cell cycle progression (Supplementary Fig. 2A). Interestingly, this TGF-β-driven transcriptional response was heavily blunted in TIF1γ-deficient cells, where only 356 genes were upregulated and 616 genes were downregulated (Supplementary Fig. 2B). These findings indicate that TIF1γ mediates a substantial component of the TGF-β transcriptional response in activated CD8⁺ T cells.

To determine whether TIF1γ possesses TGF-β-independent functions, we compared the transcriptomes of WT and TIF1γ KO cells activated in the absence of TGF-β. This comparison revealed minimal transcriptional changes, with only 29 genes upregulated and 15 downregulated in the absence of TIF1γ (Supplementary Fig. 2C). These findings indicate that TIF1γ exerts relatively little transcriptional activity in activated CD8⁺ T cells in the absence of TGF-β.

We next focused on the transcriptional landscape under TGF-β stimulation to dissect the precise programs governed by TIF1γ. Direct comparison between TGF-β-treated WT and TIF1γ KO cells revealed thousands of differentially expressed genes (DEGs) (Fig. 3A). In the presence of TGF-β, TIF1γ KO cells failed to upregulate negative regulators of activation and promoters of exhaustion (e.g., *Cd276*, *Havcr2*), cell cycle inhibitors (*Cdkn2a*, *Cdkn2b*), and the pro-apoptotic regulator *Prdm1* (Fig. 3B). Furthermore, TIF1γ deficiency blunted TGF-β-dependent induction of specific chemokine receptor genes, including *Cxcr2*, *Ccr3*, and *Xcr1*.

**Fig. 3.**
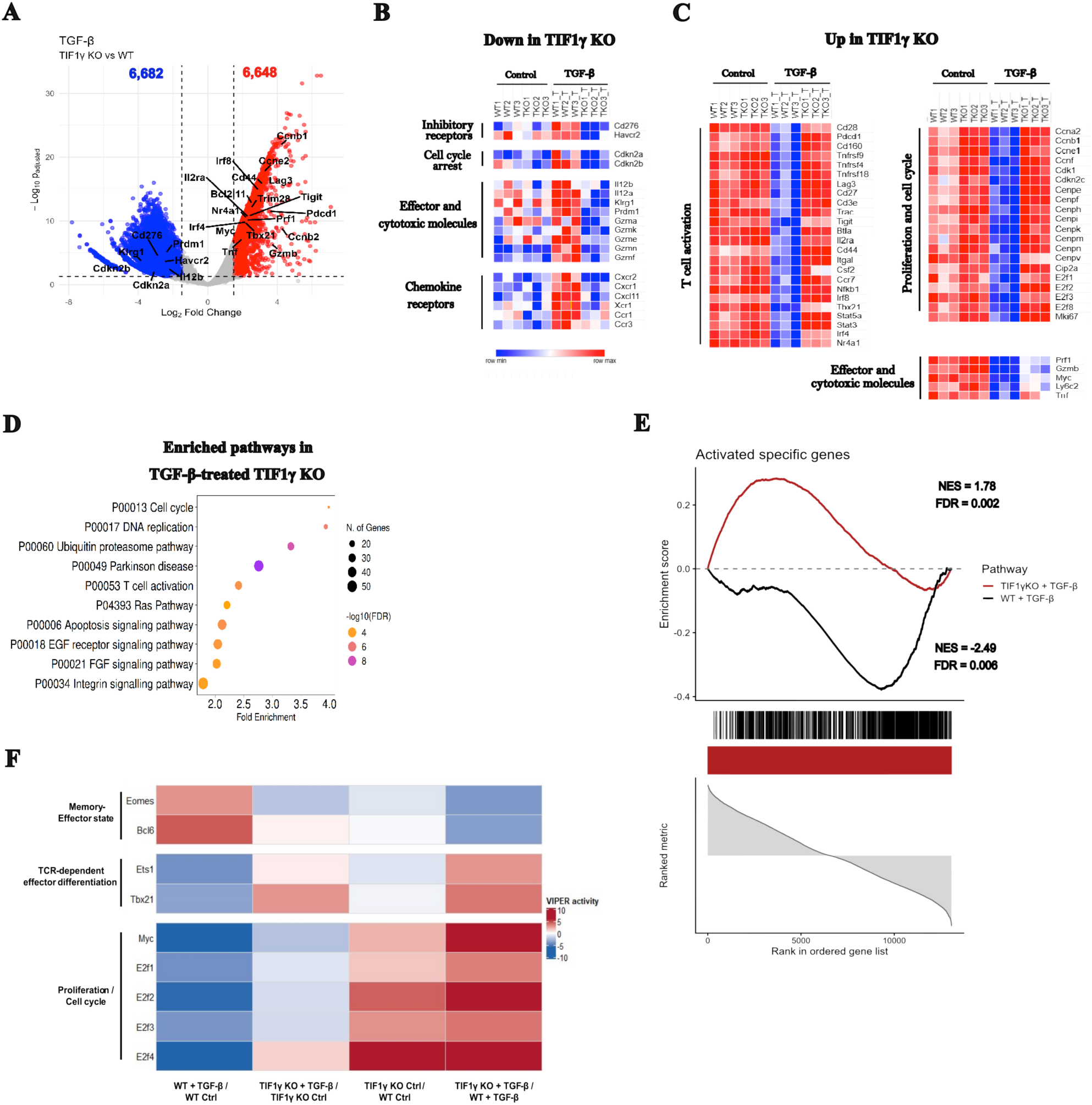
The transcriptomic landscape of TGF-β-treated TIF1γ KO CD8⁺ T cells mirrors that of activated wild-type cells in the absence of TGF-β. Naïve WT and TIF1γ KO CD8^+^ T cells were isolated and activated *in vitro* using plate-bound anti-CD3 (2 µg/mL) and anti-CD28 (1 µg/mL) for 3 days in the presence of neutralizing anti-TGF-β (2 µg/mL) or recombinant mouse TGF-β1 (2 ng/mL). (A) Volcano plot illustrating differentially expressed genes (DEGs) in recombinant TGF-β1-treated TIF1γ KO CD8^+^ T cells compared directly with TGF-β1-treated WT counterparts. (B, C) Representative lists of select, biologically relevant genes found significantly downregulated (B) or upregulated (C) in TGF-β1-treated TIF1γ KO cells relative to treated WT cells. (D) Gene Ontology (GO) biological process pathway analysis showing functional terms significantly enriched among DEGs in TGF-β1-treated TIF1γ KO versus treated WT CD8^+^ T cells. (E) Pre-ranked Gene Set Enrichment Analysis (GSEA) profiles of TGF-*β*1-treated WT and TIF1γ KO CD8^+^ T cells, evaluated against a benchmark published dataset of activated versus naïve CD8^+^ T cells^26^. NES, normalized enrichment score. Transcriptomic data were generated from RNA sequencing (RNA-seq) of 3 independent biological replicates (n = 3 mice per group). (F) Heatmap of VIPER analysis of inferred transcription factor activity.

One of the most striking observations was that genes controlling T-cell activation and cell-cycle progression remained largely resistant to TGF-β-mediated repression in TIF1γ-deficient cells. Comparative analysis highlighted a specific cluster of core T cell activation and proliferation genes that were heavily suppressed by TGF-β in WT cells but resisted this repression in TIF1γ KO counterparts (Fig. 3C). This molecular signature provides a rigorous mechanistic basis for our previous *in vitro* observations showing uncontrolled proliferation and effector differentiation in TIF1γ KO cell cultures despite TGF-β exposure (Fig. 2). In contrast, core components of the cytotoxic machinery, including *Eomes*, *Fasl*, *Ifng*, and *Il2*, were equally downregulated by TGF-β across both genotypes (Supplementary Fig. 2E, F), whereas *Prf1* and *Gzmb* experienced only partial, attenuated repression in TIF1γ KO cells (Fig. 3C). Interestingly, canonical TGF-β-induced genes (*Itgae*, *Ccr8*, *Ccr6*, and *Tgfbr1*) remained unaffected by TIF1γ deletion (Supplementary Fig. 2D), hence genes associated with cytotoxic differentiation and tissue homing remained largely responsive to TGF-β despite TIF1γ deficiency, indicating that distinct TGF-β-responsive transcriptional modules display different dependencies on TIF1γ.

To confirm the functional identity of the TIF1γ-regulated transcriptome, we performed enrichment pathway analysis, which identified cell cycle and T cell activation among the top five pathways significantly enriched in TGF-β-stimulated TIF1γ KO cells relative to TGF-β-stimulated WT controls (Fig. 3D). To define the identity of this transcriptional program further, we utilized a public dataset of the core transcriptional program that shifts during the transition from naïve to activated CD8⁺ T cells ^26^. Gene Set Enrichment Analysis (GSEA) revealed that while TGF-β-treated WT cells displayed an expected, marked inversion of the activation signature, TGF-β-treated TIF1γ KO cells retained a transcriptional profile that strongly correlated with actively proliferating, non-suppressed CD8⁺ T cells (Fig. 3E). We next used VIPER to infer transcription factor activity and define the regulatory programs associated with TIF1γ-dependent TGF-β responses. This analysis revealed a distinct TGF-β-induced transcriptional program in WT CD8⁺ T cells that was markedly attenuated or reversed in the absence of TIF1γ. Strikingly, TGF-β strongly suppressed MYC- and E2F-associated activity in WT cells, whereas this suppression was blunted in TIF1γ-deficient cells. Conversely, TGF-β increased EOMES-and BCL6-associated activity in WT cells, responses that were absent in TIF1γ-deficient cells. Notably, TGF-β also coordinately suppressed the TCR-associated transcription factors ETS1 and T-bet (Tbx21) in WT cells, whereas both activities were maintained or increased following TGF-β exposure in TIF1γ-deficient cells (Fig. 3F). Thus, TIF1γ is required to establish a specific transcriptional response to TGF-β that simultaneously represses MYC/E2F- and ETS1/T-bet-associated activation programs while promoting an EOMES/BCL6-associated differentiation program. Collectively, these findings identify TIF1γ as a selective transcriptional effector of TGF-β signaling in CD8⁺ T cells and provide a mechanistic framework linking TIF1γ-dependent TGF-β signaling to the suppression of proliferation and remodeling of T cell differentiation programs.

### TIF1γ tunes TCR signaling thresholds during TGF-β-mediated control of CD8⁺ T cells

To validate the functional relevance of our transcriptomic findings, we first examined the surface expression of key activation markers regulated by the TGF-β-TIF1γ axis. Consistent with the RNA-seq data, TIF1γ KO cells stimulated in the presence of TGF-β showed increased frequencies of CD44+, PD-1+, and CD25+ cells and elevated surface levels of CD44 and PD-1, but not CD25, compared with WT controls (Supplementary Fig. 3A-C), whereas CD69 expression remained unchanged (Supplementary Fig. 3D). We next asked whether TIF1γ specifically modulates proximal TCR-evoked signals or CD28-mediated co-stimulation.

To decouple these microenvironmental inputs, we first cultured naïve WT and TIF1γ KO cells under a fixed concentration of anti-CD28 while titrating anti-CD3 cross-linking to modulate TCR signal strength. Across a broad range of anti-CD3 concentrations, TIF1γ KO cells remained resistant to the antiproliferative effects of TGF-β (Fig. 4A, B, Supplementary Fig. 4A). This proliferative resistance was consistent with an inability of TGF-β to restrain effector differentiation (CD62L^-^ CD44^+^), even under weak TCR stimulation (Fig. 4C, D). Furthermore, TIF1γ KO cells maintained higher frequencies of CD25^+^, IRF4^+^, and Nur77^+^ populations under TGF-β exposure (Supplementary Fig. 4B–D). Interestingly, TIF1γ deficiency selectively increased IRF4 expression without altering Nur77 or CD25 levels (Supplementary Fig. 4E–G), suggesting preferential regulation of downstream transcriptional amplification rather than proximal TCR engagement.

**Fig. 4.**
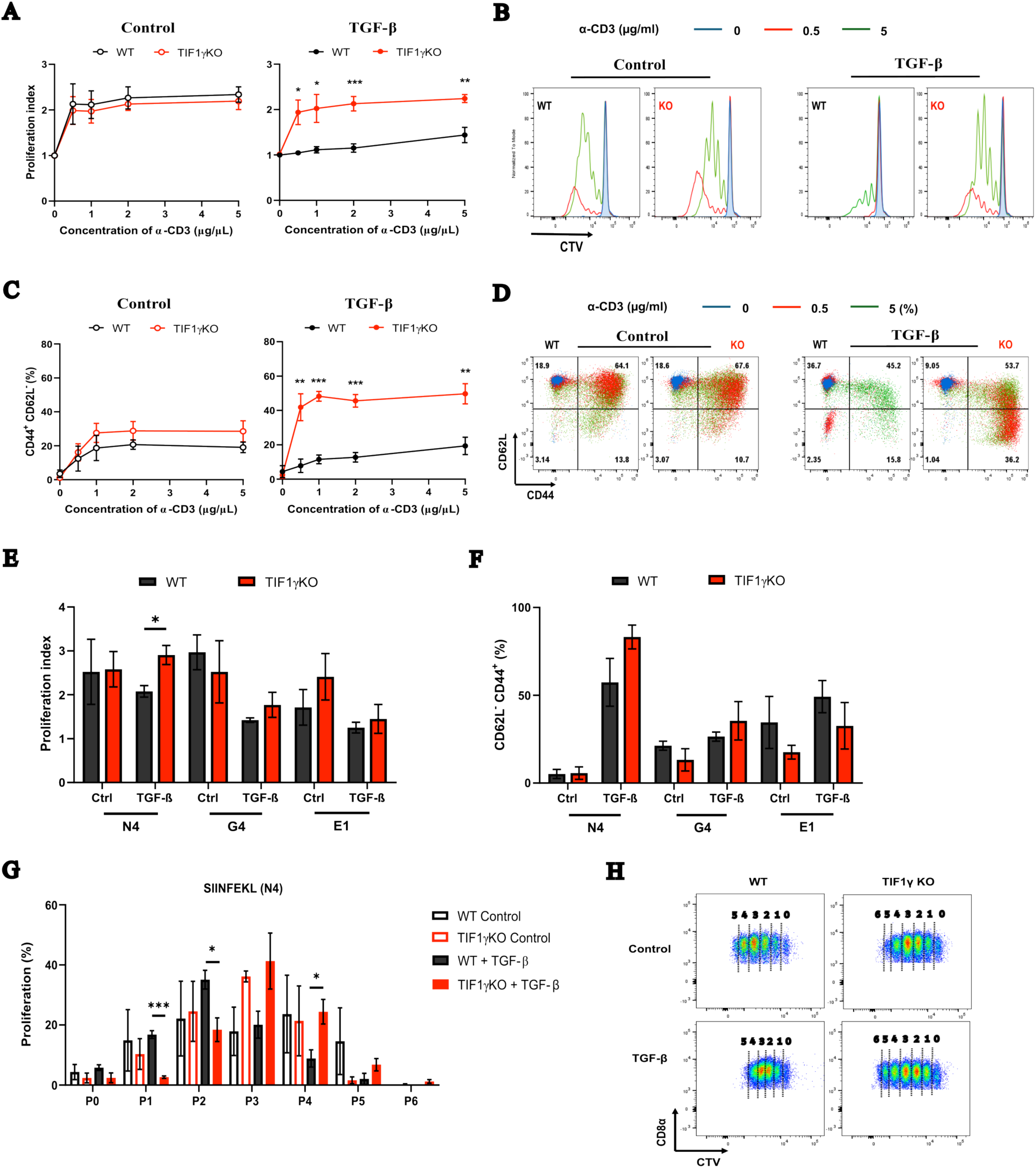
TIF1γ constraints are bypassed under strong TCR signal input or high-affinity antigen stimulation during TGF-β suppression. Naïve WT and TIF1γ KO CD8^+^ T cells were isolated and activated *in vitro* for 4 days using a fixed concentration of plate-bound anti-CD28 (1 µg/mL) and increasing concentrations of anti-CD3 (0, 0.5, 1, 2, and 5 µg/mL), in the presence of either neutralizing anti-TGF-β (2 µg/mL) or recombinant mouse TGF-β1 (2 ng/mL). (**A**) Concentration-response curves tracking the proliferation index (**B**) Representative flow cytometry histograms of CTV dilution at selected anti-CD3 concentrations: 0 *μ*g/mL (blue), 0.5 µg/mL (red), and 5 µg/mL (green). (**C**) Concentration-response curves illustrating the frequencies of the effector memory population (CD62L^-^ CD44^+^) as a function of TCR signal strength. (**D**) Representative pseudocolor dot plots of differentiation subsets identified by CD62L and CD44 gating across the indicated anti-CD3 concentrations showing selected anti-CD3 concentrations: 0 µg/mL (blue), 0.5 µg/mL (red), and 5 µg/mL (green). Panel percentages indicate values obtained at the 5 µg/mL condition. For antigen-affinity assays, naïve OT-I WT and OT-I TIF1γ KO CD8^+^ T cells were isolated from CD45.2^+^ *Tif1g^fl/fl^* and CD45.2^+^ *Cd8-Cre Tif1g^fl/fl^* mice, respectively, and co-cultured with CD45.1^+^ antigen-presenting cells (APCs) pulsed with the altered peptide ligands SIINFEKL (N4; high affinity), SIIGFEKL (G4; intermediate affinity), or EIINFEKL (E1; low affinity) for 4 days under matching anti-TGF-β or TGF-β1 conditions. (**E**) Quantification of the proliferation index (**F**) and frequencies of the effector population (CD62L^-^ CD44^+^); in response to varying TCR–peptide affinities. (**G**) Cell-division tracking of the percentage of proliferating CD8^+^ T cells within each sequential division peak following stimulation with the high-affinity N4 peptide and (**H**) representative histograms. Data represent the mean ± SEM of 3 or 4 independent experiments. Statistical significance was determined using a two-tailed unpaired Student’s t-test or two-way ANOVA for titration curves. \**P < 0.05; \**\**P < 0.01; ***P < 0.001*.

To conversely interrogate the contribution of co-stimulation, we titrated anti-CD28 concentrations against a fixed, optimal concentration of anti-CD3. In the complete absence or at 0.25 – 2 µg/mL of anti-CD28, TIF1γ KO cells consistently bypassed TGF-β-mediated growth arrest, retaining superior proliferative capacity and effector differentiation (Supplementary Fig. 5A–C). However, providing a robust, high-concentration co-stimulatory signal (4 µg/mL) rescued the WT phenotype, effectively neutralizing the advantages observed in TIF1γ KO cells (Supplementary Fig. 5A–C). Under these conditions, the frequencies of CD25^+^, IRF4^+^, and Nur77^+^ cells converged between genotypes (Supplementary Fig. 5D–G, I). However, TIF1γ KO cells showed higher levels of IRF4 in the absence and at 0.25 – 2 µg/mL of anti-CD28, losing this difference between genotypes at the higher concentration tested (Supplementary Fig. 5H). Together, these experiments demonstrate that the TGF-β-TIF1γ axis specifically restrains downstream cascades elicited by the TCR rather than modulating co-stimulatory integration, and a strong co-stimulation signal was sufficient to override the requirement for TIF1γ-mediated suppression in CD8^+^ T cells.

We next asked whether TIF1γ influences the physiological threshold of TCR discrimination against altered peptide ligands of varying affinities. To test this, we crossed TIF1γ KO mice to the OTI transgenic background and co-cultured naïve WT or TIF1γ KO OTI cells with APCs pulsed with SIINFEKL (N4, high affinity), SIIGFEKL (G4, medium affinity), or EIINFEKL (E1, low affinity) peptides ^27,28^. Upon high-affinity N4 stimulation, TIF1γ KO OTI cells exhibited accelerated proliferation compared to WT counterparts in the presence of TGF-β (Fig. 4E), and a slight but not significant increase in the activated phenotype (Fig. 4F). Tracking individual division peaks revealed that while the majority of WT cells were arrested after 1 to 2 divisions under TGF-β suppression, TIF1γ KO cells robustly progressed through 3 to 4 rounds of division (Fig. 4G, H). Interestingly, this proliferative advantage was entirely lost when cells were stimulated with the lower-affinity ligands G4 or E1 (Fig. 4G, H). These findings demonstrate that TIF1γ selectively limits proliferation driven by high-affinity TCR stimulation while having little effect on responses elicited by lower-affinity ligands. Together, these data demonstrate that TIF1γ establishes the signaling threshold above which CD8⁺ T cells escape TGF-β-mediated proliferative restraint. Because TIF1γ selectively limited responses to strong TCR stimulation in vitro, we next examined whether it similarly restrained physiological tonic TCR signaling during homeostatic proliferation in vivo.

### TIF1γ restrains homeostatic proliferation independently of the TCR repertoire

Because homeostatic proliferation is driven by low-affinity interactions with self-peptide–MHC complexes, we next asked whether TIF1γ also regulates CD8⁺ T cell responses to physiological tonic TCR signaling *in vivo* ^29–33^. To determine whether TIF1γ regulates TCR sensitivity to self-antigens in CD8⁺ T cells, we performed a competitive adoptive co-transfer assay. Naïve WT (CD45.1⁺) and naïve TIF1γ KO (CD45.2⁺) cells were labeled with CTV, mixed at a 1:1 ratio, and intravenously injected into lymphopenic *Rag1*^⁻/⁻^ recipient mice (Fig. 5A). By day 5 post-transfer, TIF1γ KO cells exhibited a marked competitive advantage, showing significantly higher frequencies and absolute numbers in the spleen compared to their WT counterparts (Fig. 5B). This accumulation correlated with a 1.7-fold increase in proliferation in TIF1γ KO cells (Fig. 5C). Phenotypic analysis revealed that while WT cells predominantly retained a naïve status (CD62L⁺ CD44⁻), a high proportion of TIF1γ KO cells rapidly differentiated into an effector-like phenotype (CD62L⁻ CD44⁺) (Fig. 5D), indicating that TIF1γ restrains homeostatic activation and differentiation under lymphopenic conditions.

**Fig. 5.**
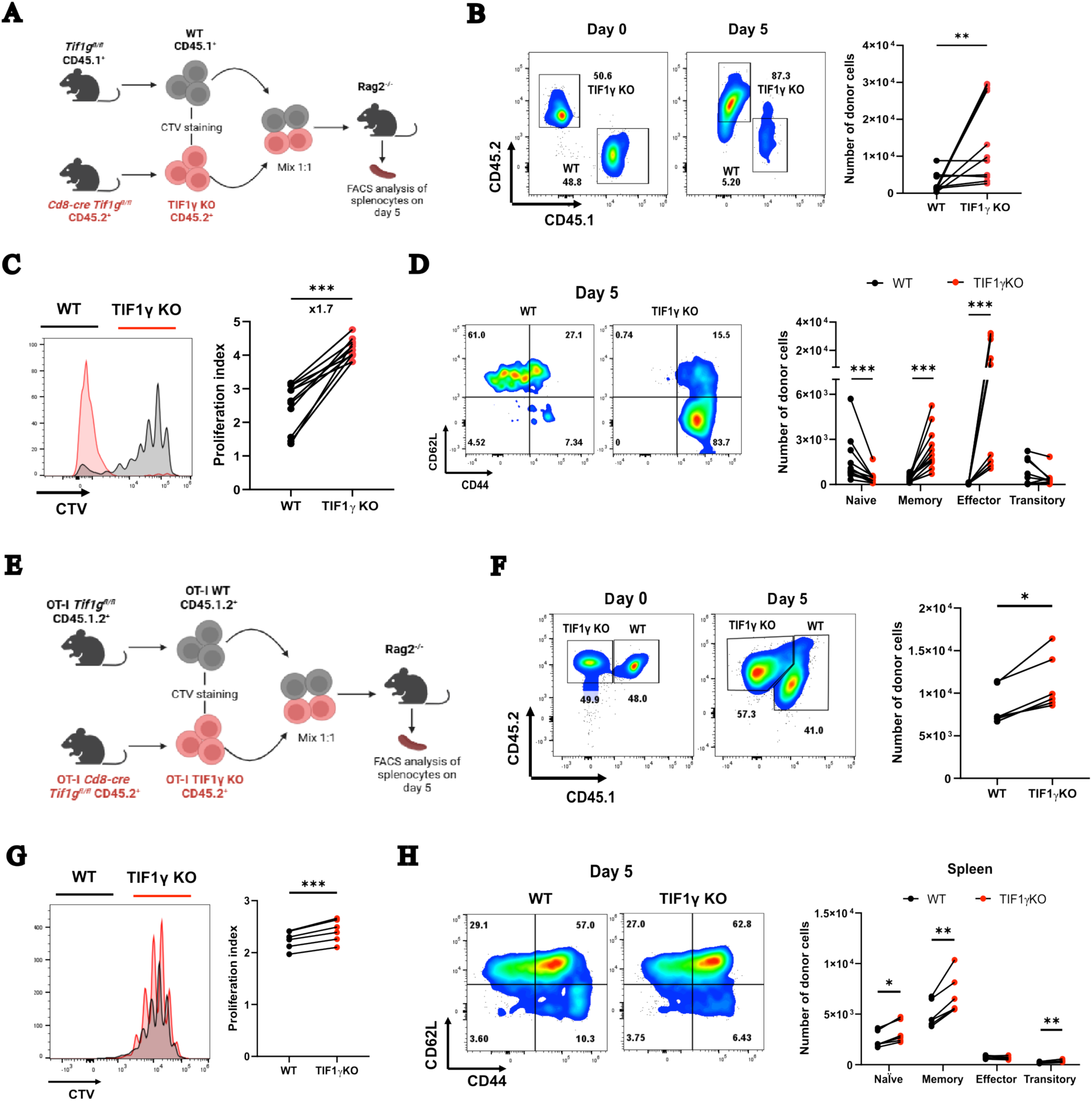
TIF1γ is essential for maintaining naïve CD8⁺ T cell quiescence and constraining lymphopenia-induced homeostatic proliferation. **(A)** Schematic overview of the polyclonal homeostatic proliferation model in lymphopenic hosts. Purified polyclonal naïve WT (CD45.1^+^) and TIF1γ KO (CD45.2^+^) CD8^+^ T cells were labeled with CTV and co-transferred at a 1:1 ratio (total 3×10^6^ cells) into Rag1^-/-^ recipient mice. (B) Representative flow cytometry plots showing the recovery frequencies at day 0 and day 5 (left) and cumulative absolute numbers (right) of donor WT and TIF1γ KO CD8^+^ T cells in the spleen at day 5 post-adoptive transfer. (**C**) Representative histograms of CTV dilution (left) and cumulative data of the proliferation index (right) of splenic donor populations. (**D**) Representative flow cytometry plots and cumulative frequencies of differentiation subsets among donor cells, classified based on CD62L and CD44 expression into naïve (CD62L^+^CD44^−^), central memory (CD62L^+^CD44^+^), effector memory (CD62L^−^CD44^+^), and transitional (CD62L^−^CD44^−^) phenotypes. (**E**) Schematic overview of the monoclonal antigen-specific homeostatic proliferation model. Naïve WT (CD45.1^+^CD45.2^+^) and TIF1γ KO (CD45.2^+^) OT-I CD8^+^ T cells were labeled with CTV and co-transferred at a 1:1 ratio into Rag1^-/-^ recipients. (**F–H**) Representative flow cytometry plots and cumulative data at day 5 post-transfer for the monoclonal OT-I model, tracking absolute donor cell numbers in the spleen (**F**), CTV dilution-derived proliferation index (**G**), and CD62L/CD44-based differentiation subset distribution (**H**). For panels A–D: data represent the mean ± SEM of three independent experiments (total n = 11 recipient mice). For panels E–H: data represent the mean ± SEM of two independent experiments (total n = 6 recipient mice). Statistical significance between co-transferred populations within the same host was determined using a two-tailed paired Student’s t-test or a Wilcoxon signed-rank test, depending on data normality. \**P < 0.05; \**\**P < 0.01; ***P < 0.001*.

To determine whether this phenotype reflected intrinsic TIF1γ deficiency rather than differences in TCR repertoire composition, we repeated the experiment using OT-I transgenic CD8⁺ T cells. Naïve OTI WT (CD45.1.2⁺) and naïve OTI TIF1γ KO (CD45.2⁺) cells were co-transferred into *Rag1*^⁻/⁻^ hosts (Fig. 5E). Consistent with our polyclonal data, OTI TIF1γ KO cells displayed augmented proliferation relative to OTI WT cells at day 5 post-transfer (Fig. 5F, G). Furthermore, despite a modest relative increase in memory-like percentages and reduced effector frequencies, the absolute numbers of naïve, memory, and transitional populations were universally elevated in TIF1γ KO OTI cells (Fig. 5H). Together, these findings demonstrate that TIF1γ intrinsically restrains homeostatic proliferation and preserves naïve CD8⁺ T cell quiescence independently of TCR repertoire diversity.

### TIF1γ selectively controls TGF-β but not γ-chain cytokine responses in CD8⁺ T cells

Because homeostatic proliferation and quiescence depend on three main signals-tonic TCR signaling, TGF-β signaling, and an IL-7 signal-we next asked whether TIF1γ specifically regulates TGF-β-mediated suppression of TCR activation or more broadly affects γ-chain cytokine signaling ^30,34,35^. To address this, WT and TIF1γ-deficient CD8⁺ T cells were stimulated with limiting anti-CD3/anti-CD28 in the presence of TGF-β and IL-7.

Exogenous TGF-β potently suppressed the proliferation of activated WT cells; however, TIF1γ KO cells were completely refractory to this inhibition (Supplementary Fig. 6A). This resistance to TGF-β-mediated suppression in TIF1γ KO cells persisted in the presence of IL-7 (Supplementary Fig. 6B, C). Phenotypically, TGF-β-treated TIF1γ KO cultures yielded an increased frequency of effector cells and a concomitant loss of the naïve pool compared to WT controls (Supplementary Fig. 6D), a skewing that remained unaffected by IL-7 supplementation (Supplementary Fig. 6E, F). Notably, these alterations were not driven by differential cell death, as apoptosis rates remained comparable between WT and TIF1γ KO cells across all conditions (Supplementary Fig. 6G, H).

To further delineate the specificity of TIF1γ to TGF-β signaling, we evaluated responses to IL-2 and IL-15—two common γ-chain cytokines that promote T cell survival and robust proliferation. Consistent with earlier observations, IL-2 restored proliferative responses in WT cells but failed to normalize effector differentiation in TIF1γ-deficient cultures, whereas IL-15 had little effect on either process (Supplementary Fig. 6I-N). Together, these findings demonstrate that TIF1γ selectively mediates TGF-β-dependent regulation of TCR-driven activation while leaving γ-chain cytokine responses largely intact.

### TIF1γ is dispensable during primary infection but required for controlled CD8⁺ T cell recall responses

Having established that TIF1γ maintains homeostatic quiescence, we investigated whether this alternative TGF-β branch is required to restrain CD8⁺ T cell activation and clonal expansion under acute, highly inflammatory settings *in vivo*. We first challenged WT and TIF1γ KO mice with ovalbumin-expressing *Listeria monocytogenes* (*Lm*-OVA) and monitored the longitudinal primary immune response in peripheral blood (Supplementary Fig. 7A). The frequencies, absolute numbers, and differentiation kinetics of endogenous antigen-specific (Tetramer^+^) and nonantigen-specific (Tetramer^-^) CD8⁺ T cells remained entirely comparable between genotypes in the circulation, as well as within the spleen and liver at the peak of clonal expansion on day 7 (Supplementary Fig. 7B–G, Supplementary Fig. 8A–C). These findings indicate that the strong inflammatory environment associated with primary bacterial infection can compensate for the absence of TIF1γ during initial CD8⁺ T cell expansion.

Interestingly, a closer examination of the non-antigen-specific CD8⁺ T cell pool revealed a localized phenotypic aberration: TIF1γ KO mice displayed a significant accumulation of terminally differentiated long-lived effector cells (LLEC; KLRG1^+^ CD127^+^) within the spleen and liver on day 7 (Supplementary Fig. 8D–F). However, this phenotypic difference did not translate into global differences in the production of effector/cytolytic molecules in CD8^+^ T cells between genotypes (Supplementary Fig. 7H–M, Supplementary Fig. 8G–L). These findings revealed a selective alteration in bystander CD8⁺ T cell differentiation during acute inflammation, prompting us to examine whether TIF1γ deficiency affected subsequent recall responses.

To test this, we reinfected mice with *Lm*-OVA 30 days after the primary challenge (Fig. 6A). In contrast to the primary response, TIF1γ KO mice displayed marked expansion of antigen-specific, but not nonantigen-specific, memory CD8⁺ T cells within the blood, spleen, and liver by day 3 post-reinfection (Fig. 6B–D, Supplementary Fig. 9A–C), establishing a non-redundant role for TIF1γ in restraining secondary clonal expansion. Phenotypic dissection at this early recall phase revealed that TIF1γ KO antigen-specific compartments displayed altered differentiation dynamics characterized by increased MPEC and LLEC populations with reduced EEC representation (Fig. 6E–G). Concurrently, the non-antigen-specific LLEC accumulation observed during primary infection persisted through the recall phase in the spleen and liver of challenged mice (Supplementary Fig. 9D–F). Together, these findings reveal that TIF1γ functions as a context-dependent regulator of CD8⁺ T cell responses. While strong inflammatory signals during primary infection compensate for TIF1γ deficiency, this pathway becomes essential during recall responses to restrain excessive expansion and maintain appropriate memory differentiation.

**Fig. 6.**
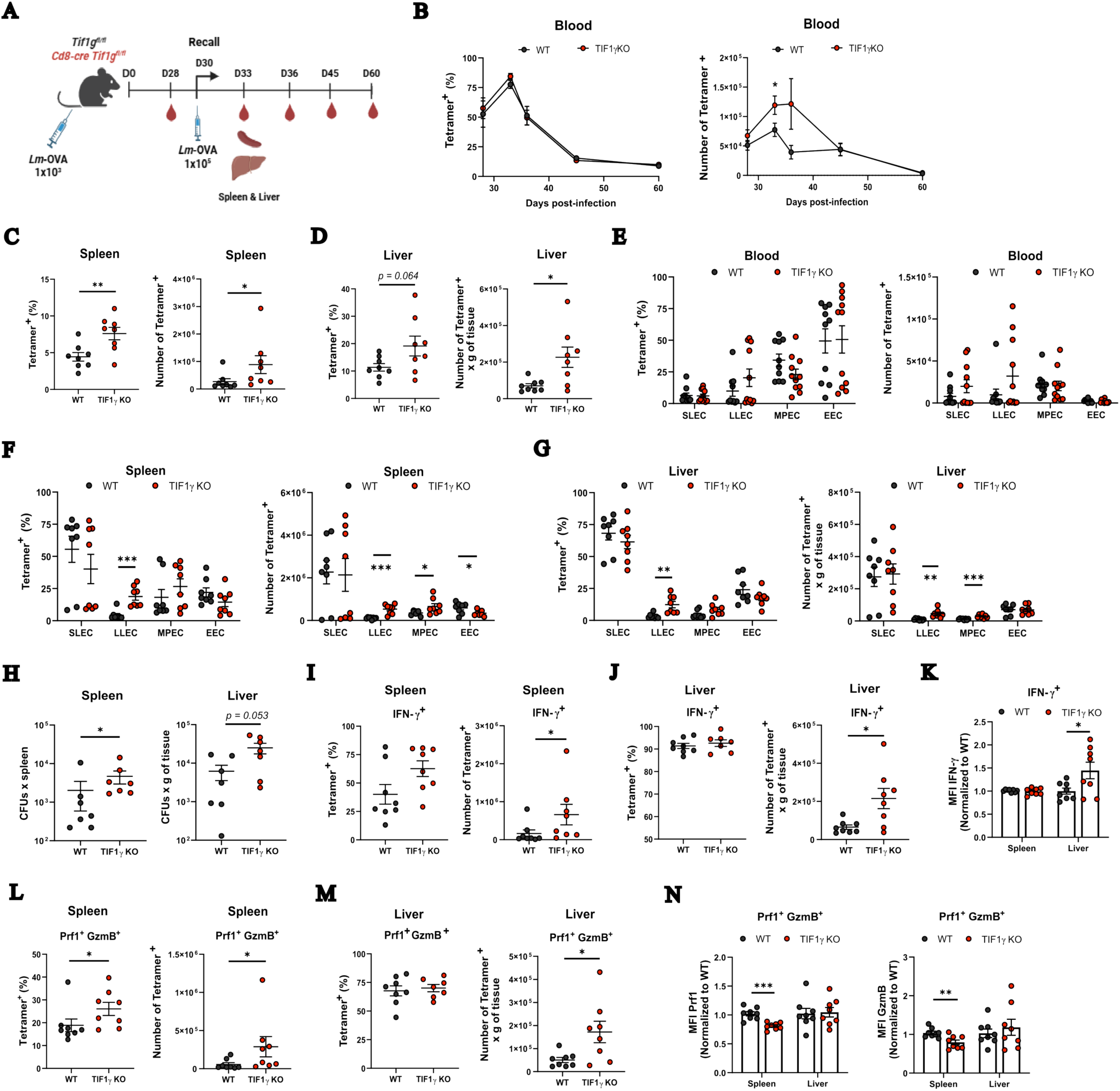
Massive expansion but defective cytotoxicity of TIF1γ KO CD8⁺ T cells upon secondary *Lm*-OVA challenge. (**A**) Schematic experimental timeline of the secondary challenge (rechallenge) model. WT and TIF1γ KO mice were primed with 1×10^3^ CFUs of *Lm*-OVA. At day 30 post-infection, mice were reinfected with 1×10^5^ CFUs of *Lm*-OVA; longitudinal peripheral blood was sampled on days –28, 33, 36, 45, and 60, and spleens and livers were harvested at day 33 (day 3 post-rechallenge). (**B**) Kinetic curves showing the frequencies and absolute numbers of antigen-specific (OVA-tetramer^+^) CD8^+^ T cells in peripheral blood. (**C, D**) Frequencies and absolute numbers of OVA-tetramer^+^ CD8^+^ T cells in the spleen (**C**) and liver (**D**) at day 3 post-rechallenge. (**E–G**) Frequencies and absolute numbers of antigen-specific CD8^+^ T cell memory subsets, differentiated into short-lived (SLECs; KLRG1^+^CD127^−^), long-lived effector cells (LLEC; KLRG1^+^CD127^+^), memory precursor effector cells (MPEC; KLRG1^−^CD127^+^), and early effector cells (EEC; KLRG1^−^CD127^−^), analyzed in peripheral blood (**E**), spleen (**F**), and liver (**G**) at day 3 post-rechallenge (day 33 post-prime). (**H**) Bacterial burdens (CFUs) quantified in the spleen (left) and liver (right) 60 hours post-reinfection in WT and TIF1γ KO mice challenged with 5×10^5^ CFUs of *Lm*-OVA at day 30 post-primary infection. (**I, J**) Percentages and absolute numbers of IFN-γ^+^ OVA-tetramer^+^ CD8^+^ T cells in the spleen (**I**) and liver (**J**) at day 3 post-rechallenge. (**K**) MFI of IFN-γ in splenic and hepatic antigen-specific CD8^+^ T cells. (**L, M**) Percentages and absolute numbers of Prf1^+^ GzmB^+^ OVA-tetramer^+^ CD8^+^ T cells in the spleen (**L**) and liver (**M**) at day 3 post-rechallenge. (**N**) MFI of Prf1 and GzmB in splenic and hepatic antigen-specific CD8^+^ T cells. All MFI values are presented normalized to their respective WT counterparts. Data represent the mean ± SEM of three independent experiments. For panels B and E: WT, n = 10; TIF1γ KO, n = 11. For panel H: WT, n = 7; TIF1γ KO, n = 7. For panels C, D, and I–N: WT, n = 8; TIF1γ KO, n = 8. Statistical significance was determined using a two-tailed unpaired Student’s t-test or a Mann-Whitney U test, depending on data normality, or a one-sample t-test for normalized MFI values. \**P < 0.05; \**\**P < 0.01; ***P < 0.001*.

### Hyper-expansion of recalled TIF1γ-deficient CD8⁺ T cells uncouples numerical abundance from protective cytotoxic immunity

Because TIF1γ KO mice exhibited increased expansion of antigen-specific CD8⁺ T cells, we asked whether this hyper-expansion translated into enhanced protective immunity. We quantified bacterial burden by enumerating colony-forming units (CFUs) in the spleen and liver at 60 hours post-reinfection. Interestingly, despite the increased antigen-specific CD8⁺ T cell expansion, TIF1γ-deficient mice exhibited impaired pathogen clearance, with significantly higher bacterial burdens in both spleen and liver (Fig. 6H).

To resolve this paradox, we characterized the quantitative and qualitative functional status of the recalled effectors. Numerically, TIF1γ KO mice possessed significantly higher absolute numbers of IFN-γ^+^ and Prf1^+^ GzmB^+^ antigen-specific CD8⁺ T cells (Fig. 6I–M). Furthermore, TIF1γ KO mice accumulated higher frequencies and absolute numbers of non-antigen-specific cytotoxic T cells in both the spleen and liver (Supplementary Fig. 9G–K). Thus, impaired bacterial control was not due to insufficient generation of antigen-specific effector cells. However, on a per-cell basis, TIF1γ-deficient CD8⁺ T cells showed markedly reduced Prf1 and GzmB expression within the tissue parenchyma in both antigen-specific and non-antigen-specific cell subsets (Fig. 6N, Supplementary Fig. 9L). This uncoupling demonstrates that TIF1γ-deficient cells readily acquire effector phenotypes but fail to achieve equivalent cytotoxic competency. These *in vivo* observations align with our transcriptomic and in vitro data, showing that the alternative TIF1γ branch limits activation and expansion while also acting as a molecular signal-threshold regulator to sustain cytotoxic efficiency. Thus, our findings reveal a fundamental regulatory principle of CD8⁺ T cell immunity: TGF-β signaling through TIF1γ limits excessive clonal expansion while preserving the cytotoxic capacity required for pathogen control.

Finally, we asked whether this functional impairment extended to a tumor setting. Following B16-OVA engraftment, TIF1γ KO mice exhibited significantly accelerated tumor growth and larger tumor volumes compared to WT controls (Supplementary Fig. 10A, B). This failure to control the malignancy correlated with a marked reduction in the frequencies of both antigen-specific and bystander Prf1^+^ GzmB^+^ tumor-infiltrating lymphocytes (TILs) (Supplementary Fig. 10C, F). Moreover, TIF1γ-deficient TILs exhibited a selective, significant decrease in Prf1 expression levels compared to WT counterparts (Supplementary Fig. 10D, E, H, I). Collectively, these findings demonstrate that increased CD8⁺ T cell abundance is insufficient to ensure protective immunity when cytotoxic competency is compromised. Across infection and tumor models, TIF1γ preserves the functional quality of CD8⁺ T cells by coordinating the balance between expansion and effector execution.

## Discussion

Naïve CD8⁺ T cells must maintain quiescence while preserving the capacity to rapidly proliferate and acquire effector functions upon antigen encounter. How these opposing requirements are coordinated remains a fundamental question in T cell biology. Although TGF-β contributes to maintaining CD8⁺ T cell quiescence ^36^, the mechanisms by which it restrains proliferation without compromising functional competence remain incompletely understood. Here, we identify TIF1γ as a critical component of this regulatory axis. Rather than functioning as a general suppressor of CD8⁺ T cell activation, TIF1γ selectively constrains activation, proliferation, and differentiation in a context-dependent manner while contributing to the preservation of cytotoxic competence. Consistent with this model, TIF1γ was largely dispensable during the strong inflammatory environment of primary bacterial infection but became essential under conditions in which CD8⁺ T cells depend on tighter control of activation, including homeostatic proliferation, recall responses, and tumor immunity.

A central challenge in defining this pathway was the ability of strong cytokine and costimulatory signals to mask the inhibitory effects of TGF-β. In activated CD8⁺ T cell cultures, IL-2 efficiently opposed the antiproliferative and pro-apoptotic effects of TGF-β in TIF1γ-sufficient cells, illustrating how inflammatory and growth-promoting signals can override TIF1γ-dependent restraint. However, this rescue was not uniform across effector functions. Although IL-2 restored IFN-γ and GzmB expression in TIF1γ-deficient cells, it failed to restore Prf1 expression. These observations underscore that individual components of the cytotoxic program are governed by distinct regulatory mechanisms and further support the concept that proliferative expansion and cytotoxic competence can be uncoupled ^3^. Thus, the requirement for TIF1γ is likely to become most apparent in settings in which strong IL-2 signals might be limited, such as homeostatic proliferation, recall responses, and tumors with strong immunoregulatory activity. In these contexts, loss of TIF1γ permits excessive proliferation and effector differentiation while failing to preserve the full cytotoxic program.

Our transcriptional and phenotypic analyses provide evidence for such a functional uncoupling. In the absence of TIF1γ, CD8⁺ T cells readily acquire an activated effector phenotype, yet fail to establish the complete transcriptional and protein expression programs required for cytolytic function. TIF1γ-deficient mice accumulated increased numbers of antigen-specific CD8⁺ T cells in target tissues despite markedly reduced expression of Prf1 and GzmB. These findings argue against a model in which TIF1γ simply suppresses CD8⁺ T cell activation. Instead, TIF1γ appears to impose a quantitative constraint on activation that limits excessive expansion and differentiation while allowing the acquisition and maintenance of functional competence. Consistent with this interpretation, activated TIF1γ-deficient CD8⁺ T cells exhibited a markedly attenuated transcriptional response to TGF-β despite preserved TGFβRI expression, indicating that TIF1γ contributes to intracellular responsiveness to TGF-β rather than regulating receptor availability. Importantly, the loss of TIF1γ preferentially altered activation- and proliferation-associated transcriptional programs while leaving substantial portions of the canonical TGF-β transcriptional response intact. This selective effect is consistent with previous studies indicating that Smad4-dependent TGF-β signaling preferentially regulates tissue localization, migration, and related transcriptional programs ^9,11,13^, and supports a model in which TGF-β signaling is partitioned into functionally distinct transcriptional modules.

The context dependence of this pathway further suggests that TIF1γ acts as a regulator of activation thresholds rather than as an obligatory brake on CD8⁺ T cell responses. During primary *Listeria monocytogenes* infection, the strong inflammatory and costimulatory environment largely bypassed the requirement for TIF1γ. In contrast, TIF1γ became critical during homeostatic proliferation and secondary challenge, when CD8⁺ T cells encounter substantially different signaling environments. Memory CD8⁺ T cells are intrinsically poised to undergo rapid proliferation upon antigen re-encounter ^37^, and loss of TIF1γ further increased their expansion and accelerated differentiation. TCR signal strength is a well-established determinant of CD8⁺ T cell proliferation, differentiation, and memory formation ^38,39^. Our findings place TIF1γ within this regulatory framework by suggesting that TGF-β, through TIF1γ, sets the threshold at which TCR-driven activation is restrained. Thus, rather than preventing activation altogether, TIF1γ selectively limits responses once TCR stimulation exceeds a critical threshold, thereby allowing productive immune responses while preventing excessive expansion and loss of functional competence.

This threshold-setting function also extends to non-antigen-specific CD8⁺ T cells during systemic inflammation. TIF1γ-deficient non-antigen-specific CD8⁺ T cells differentiated toward an LLEC phenotype during acute infection despite lacking cognate antigen stimulation, indicating that TIF1γ normally limits inflammation-driven activation in cells that have not received a productive TCR signal. These findings suggest that TIF1γ helps preserve a reservoir of quiescent CD8⁺ T cells during systemic inflammation by restricting cytokine-driven activation in the absence of cognate antigen. Intriguingly, public transcriptomic datasets indicate that *Tif1g* (*Trim33*) expression decreases following TCR activation, whereas our data show that TGF-β increases *Tif1g* expression ^40^. Together, these observations suggest that TIF1γ may represent a dynamically regulated component of the T cell signaling network: TCR activation reduces TIF1γ expression to permit robust responses, whereas TGF-β counterbalances this process by increasing TIF1γ expression. Such reciprocal regulation provides a potential mechanism for integrating activating and suppressive signals within the same cell and may allow T cells to transition between quiescent and activated states without completely losing responsiveness to environmental cues.

The distinct phenotypes associated with disruption of TGF-β receptors, Smad4, and TIF1γ further support functional partitioning within the TGF-β signaling network. T cell-specific deletion of TGFβRII results in severe systemic immune dysregulation and uncontrolled homeostatic expansion ^30,36^, whereas Smad4 deficiency produces a substantially different phenotype characterized prominently by chronic intestinal inflammation ^11^. These divergent outcomes argue against a model in which all TGF-β signals converge on a single transcriptional program. Instead, our findings support a model in which canonical and TIF1γ-dependent pathways regulate complementary aspects of CD8⁺ T cell biology. Previous work has implicated Smad4-dependent signaling in tissue localization, migration, and regulation of cytotoxic differentiation ^9,11,13^, whereas our data identify TIF1γ as a major regulator of activation threshold, proliferation, and quiescence. Importantly, these pathways should not be viewed as completely independent. Rather, their functional interaction may allow TGF-β to coordinate the magnitude of CD8⁺ T cell expansion with the acquisition of appropriate effector function.

Our transcription factor activity analysis provides a clear mechanistic framework for this phenotype. In WT CD8^+^ T cells, TGF-β strongly suppressed MYC- and E2F-driven transcriptional networks, enforcing a non-proliferative state. In contrast, TIF1γ-deficient cells failed to enact this anti-proliferative program, maintaining active MYC and E2F signaling despite TGF-β exposure. At the same time, while TGF-β induced EOMES and BCL6 transcriptional activity in WT cells, this response was completely lost in the absence of TIF1γ. Given the established roles of EOMES and BCL6 in driving memory differentiation ^41,42^ and sustaining stem-like progenitor T cells, the loss of TIF1γ shifts CD8^+^ T cells away from a memory fate and toward terminal effector differentiation. Indeed, TGF-β/Smad2 signaling normally induces BCL6 to prevent BLIMP-1-mediated terminal exhaustion ^43^; consistent with this, TIF1γ-deficient cells exposed to TGF-β showed abnormally high PD-1 levels. Together, these data demonstrate that TIF1γ is required for TGF-β to halt the cell cycle and direct memory lineage specification, explaining why TIF1γ-deficient CD8^+^ T cells fail to control tumor growth.

Notably, TGF-β-dependent suppression of ETS1 and T-bet-associated activity was also lost or reversed following TIF1γ deficiency. Given the established roles of ETS1 and T-bet in integrating TCR-dependent signals and regulating CD8⁺ T cell differentiation, these changes raise the possibility that TIF1γ influences the transcriptional interpretation of TCR signals downstream of TGF-β. Importantly, these VIPER-derived activities represent inferred regulatory states rather than direct measurements of transcription factor occupancy or protein activity. However, the coordinated changes in MYC/E2F, EOMES/BCL6, and ETS1/T-bet activity provide a coherent transcriptional framework linking TIF1γ to the regulation of proliferation and differentiation.

Mechanistically, these findings extend previous models of TGF-β-mediated repression of CD8⁺ T cell effector function. Early studies demonstrated that TGF-β induces CREB and ATF1 to cooperate with Smad2 and Smad3 in repressing *Ifng* and *Gzmb* ^3^, whereas subsequent work established an essential role for Smad4 in the suppression of GzmB and T-bet in TGF-β-stimulated CD8⁺ T cells ^11^. However, the consequences of Smad4-dependent signaling are strongly influenced by cellular and inflammatory context. Indeed, Smad4-deficient CD8⁺ T cells can exhibit reduced cytotoxic gene expression, but not impaired expansion, during acute *Lm*-OVA and X31-OVA infections ^12,13^, or within tumors ^14^, highlighting Smad4 as a positive regulator of the activated cytotoxic program. Our findings expand this paradigm by introducing TIF1γ as a non-redundant partner in this network. Although TIF1γ-deficient CD8⁺ T cells are resistant to the antiproliferative effects of TGF-β, they suffer from a severe reduction in Prf1 and GzmB levels *in vivo* under both resting and inflammatory conditions. The TIF1γ-dependent pathway primarily restrains activation and preserves cellular quiescence, whereas cooperation between TIF1γ and the canonical Smad4 pathway coordinates cytotoxic differentiation. More broadly, our findings demonstrate that effective CD8⁺ T cell immunity depends not simply on the magnitude of clonal expansion but on the coordinated regulation of expansion and functional quality. By establishing the signaling threshold through which TGF-β calibrates these processes, TIF1γ emerges as a central regulator of protective cellular immunity.

Collectively, our findings support a model in which TGF-β signaling is functionally partitioned into complementary branches rather than acting as a uniform suppressive pathway. Within this framework, the TIF1γ-dependent branch establishes the activation threshold that restrains excessive clonal expansion while preserving cellular quiescence, whereas cooperation between TIF1γ and the canonical Smad4 pathway coordinates cytotoxic differentiation. This organization allows TGF-β to independently regulate activation, proliferation, and effector function, thereby ensuring that protective immunity depends on both the magnitude and the quality of the CD8⁺ T cell response.

## Materials and Methods

### Mice

Mice were generated and maintained under specific pathogen-free conditions at the core animal facility of the Instituto de Fisiología Celular (IFC), Universidad Nacional Autónoma de México, in accordance with institutional guidelines approved by the Institutional Animal Care and Use Committee (IACUC-IFC, protocol PLL-220-23) and the CICUAL-INCMNSZ (protocol IRE-2092). All animals were on a C57BL/6 background and expressed the CD45.2 congenic marker unless otherwise indicated. To generate *Cd8a-Cre Tif1g^fl/fl^* conditional knockout mice, *Tif1g^fl/fl^* mice (kindly provided by Dr. V. Kaartinen) were crossed with C57BL/6-Tg(*Cd8a-cre*)1Itan/J (E8I-Cre) mice. C57BL/6-Tg(*TraTcrb*)1100Mjb/J (OT-I) and B6.129S7-Rag1^tm1Mom^/J (Rag1^-/-^) strains were originally purchased from The Jackson Laboratory. Experiments were conducted using age- and sex-matched mice between 8 and 14 weeks of age. Both male and female mice were included across all cohorts, and no sex-dependent differences were observed. Unless otherwise specified, experimental groups were age- and littermate-matched.

### CD8^+^ T cell isolation and culture

Naïve CD8^+^ T cells (CD62L^+^ CD44^-^) were isolated from single-cell suspensions using the EasySep Mouse Naïve CD8^+^ T Cell Isolation Kit (STEMCELL Technologies) according to the manufacturer’s instructions. Purified naïve CD8^+^ T cells (2×10^5^ cells per well) were seeded in 96-well flat-bottom tissue culture plates and maintained in complete RP10 medium (RPMI 1640 supplemented with 10% heat-inactivated FBS, 2 mM L-glutamine, 100 U/mL penicillin-streptomycin, 1 mM sodium pyruvate, 0.05 mM β-mercaptoethanol, and 1 mM HEPES). For polyclonal activation, plates were pre-coated with anti-CD3 (clone 145-2C11; Tonbo Biosciences) and anti-CD28 (clone 37.51; BioLegend) antibodies at the concentrations specified for each experiment. For antigen-specific stimulation, CD45.2^+^ naïve OT-I T cells were co-cultured with CD45.1^+^ splenic antigen-presenting cells (APCs) pulsed with 1 µM of SIINFEKL (N4), SIIGFEKL (G4), or EIINFEKL (E1) peptides. The G4 and E1 altered peptide ligands were kindly provided by Dr. José C. Crispín. Where indicated, cultures were supplemented with neutralizing anti-TGF-β1,2,3 antibody (2 µg/mL; clone 1D11.16.8, Bio X Cell) or recombinant mouse TGF-β1 (2 ng/mL; R&D Systems). Recombinant human IL-2 (20 ng/mL; PeproTech) was added when specified. All cultures were incubated at 37°C in a humidified incubator with 5% CO_2_ for 96 hours.

### Flow cytometry

Detailed information regarding antibodies, fluorochromes, and specific staining conditions is provided in the Supplementary Information (Supplementary Tables 1-2). Cell viability was assessed using the LIVE/DEAD Fixable Aqua Dead Cell Stain Kit (Thermo Fisher Scientific) in 1X PBS. For surface marker staining, cells were incubated with antibodies diluted in FACS buffer (1X PBS supplemented with 2% FBS). For intranuclear transcription factor staining, cells were fixed and permeabilized using the Foxp3/Transcription Factor Staining Buffer Set (Thermo Fisher Scientific) according to the manufacturer’s protocol. For intracellular cytokine staining (ICS), cells were fixed with 4% paraformaldehyde and permeabilized using the BD Cytofix/Cytoperm kit (BD Biosciences). For apoptosis assays, cells were resuspended and stained in Annexin V Binding Buffer (BioLegend). Samples were acquired on either an Attune NxT Flow Cytometer (Thermo Fisher Scientific) or a Cytek Aurora Spectral Flow Cytometer (Cytek Biosciences).

### RNA extraction and Quantitative Real-Time PCR (RT-qPCR)

Total RNA was extracted from sorted CD4^+^, CD8^+^, and Treg cell populations, or *in vitro*-activated CD8^+^ T cells, using TRIzol reagent (Invitrogen) following the manufacturer’s protocol. Complementary DNA (cDNA) was synthesized using SuperScript II Reverse Transcriptase (Invitrogen) according to the manufacturer’s instructions. Quantitative PCR was performed using KAPA SYBR FAST qPCR Master Mix (1x; Kapa Biosystems). Relative mRNA expression of *Tif1g* was normalized to *Actb* (β-actin) as an internal reference gene using the 2^“##*C*T^ method. All reactions were performed in technical duplicates across independent biological experiments. The specific primer sequences for mouse *Tif1g* were as follows: Forward: 5′-GGAGCTGTGAAACAAGAGAAG-3′. Reverse: 5′-AAGGTTAGTTGAAAGAGGTGG-3′.

#### In vitro cytotoxicity assays

Naïve WT and TIF1γ KO OT-I T cells were activated *in vitro* as described above and cultured in the presence of either neutralizing anti-TGF-β1,2,3 (2 µg/mL) or recombinant mouse TGF-β1 (2 ng/mL) for 72 hours. Following incubation, viable T cells were enumerated and co-cultured with B16-OVA melanoma target cells at a defined effector-to-target (E:T) ratio of 10:1 in 96-well round-bottom plates. After 6 hours of co-culture, plates were centrifuged, and supernatants were collected to quantify extracellular lactate dehydrogenase (LDH) release using the CyQUANT LDH Cytotoxicity Assay Kit (Thermo Fisher Scientific) according to the manufacturer’s instructions. Absorbance was measured at 490 nm and 680 nm using a microplate reader to calculate specific lysis percentage. In parallel, cells were harvested and analyzed by flow cytometry to determine the absolute number, phenotype, and viability of the remaining OT-I cells.

#### Adoptive cell transfers and *Listeria monocytogenes* infections

For polyclonal homeostatic proliferation assays, naïve CD8^+^ T cells were isolated from CD45.1^+^ WT and CD45.2^+^ TIF1γ KO mice, labeled with CellTrace Violet (CTV; Thermo Fisher Scientific), and co-transferred at a 1:1 ratio (total of 1×10^6^ cells) intravenously into recipient Rag1^-/-^ mice. Recipient mice were euthanized on day 5 post-transfer, and spleens were harvested for flow cytometric analysis of the transferred populations. For antigen-specific homeostatic proliferation assays, naïve OT-I T cells were isolated from CD45.1.2^+^ OT-I WT and CD45.2^+^ OT-I TIF1γ KO mice, labeled with CTV, and co-transferred at a 1:1 ratio (total of 1×10^6^ cells) into Rag1^-/-^ recipients, followed by splenocyte harvest and analysis at day 5.

For the primary infection model, WT and TIF1γ KO mice were infected intravenously via the tail vein with 1×10^3^ CFUs of *Listeria monocytogenes*-OVA (*Lm*-OVA). Longitudinal peripheral blood samples (100 µL per mouse) were collected retro-orbitally or via the submandibular vein 1 day prior to infection, and sequentially on days 5, 7, 10, 15, and 28 post-infection. In parallel cohorts, mice were euthanized on day 7 post-infection to harvest spleens and livers for immunophenotypic analysis. For secondary challenge experiments, mice were intravenously reinfected on day 30 post-primary infection with 1×10^5^ CFUs of *Lm*-OVA. Peripheral blood was sampled 2 days prior to challenge (day 28 post-primary infection) and on days 33, 36, 45, and 60 after the initial infection. In separate experimental cohorts, mice were euthanized 3 days post-reinfection (day 33 total), and spleens and livers were harvested to assess secondary CD8^+^ T cell responses via flow cytometry.

### CFU determination in the Lm-OVA infection model

For secondary challenge experiments, wild-type (WT) and TIF1γ KO mice previously infected with 1×10^3^ colony-forming units (CFUs) of *Lm*-OVA were reinfected intravenously with 5×10^5^ CFUs 30 days after the primary infection. Mice were euthanized 60 hours post-reinfection to determine bacterial burden. Spleens and livers were harvested and individually homogenized in sterile 1X PBS containing 0.1% Triton X-100. Serial dilutions of the tissue homogenates were plated onto Brain Heart Infusion (BHI) agar plates. Bacterial CFUs were enumerated after 24 to 48 hours of incubation at 37°C.

### In vivo killing assays

WT and TIF1γ KO mice were immunized subcutaneously with 100 µg of OVA protein (InvivoGen) emulsified in 50 µg of complete Freund’s adjuvant (InvivoGen). On day 21 post-immunization, target splenocytes were isolated from naïve syngeneic mice and either pulsed with the OVA_257-264_ peptide (SIINFEKL; Sigma-Aldrich, S7951-1MG) or left unpulsed. Unpulsed control splenocytes were labeled with 2.5 µM CTV (CTV^high^), whereas peptide-pulsed splenocytes were labeled with 0.25 µM CTV (CTV^low^). Labeled cells were counted, mixed at a 1:1 ratio (confirmed by flow cytometry) in 1× PBS, and a total of 3 - 4 × 10^6^ cells were intravenously injected into immunized or naïve control recipient mice. After 12–16 h, spleens were harvested, and cell suspensions were analyzed by flow cytometry. The percentage of specific lysis was calculated using the following formula: [1-(%CTV^low^ naïve / %CTV^high^ naïve) / (%CTV^low^ immunized / %CTV^high^ immunized)] x 100.

#### Melanoma mouse model

B16-OVA cells were thawed and cultured in DMEM supplemented with 10% FBS, 100 U/mL penicillin–streptomycin, and 1 mM non-essential amino acids. Cells were maintained under standard conditions and harvested upon reaching 70–80% confluence. Subsequently, 2 × 10^5^ cells were subcutaneously injected into the right flank of recipient mice. Tumor growth was monitored, and measurements were recorded up to day 21 post-inoculation. Tumor volume was calculated with the following equation: *V_T_* = (*Larger diameter* ∗ (*Smaller diameter*)^2^) ∗ 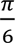. All mice were euthanized on day 21, and tumors, as well as draining and non-draining lymph nodes, were collected and processed for flow cytometry analysis of CD8^+^ T cells. To enrich immune cell populations from tumor tissue, a density gradient centrifugation using OptiPrep™ (Cat. No. D1556, Sigma-Aldrich) was performed.

#### RNA-sequencing and informatics analysis

Bulk RNA sequencing (RNA-seq) was performed on naïve WT and TIF1γ KO cells following activation with anti-CD3 (2 µg/mL) and anti-CD28 (1 µg/mL) antibodies under two conditions: in the presence of 2 µg/mL anti-TGF-β1,2,3 or 2 ng/mL recombinant TGF-β1 for 72 h. A total of six samples per genotype were analyzed, comprising three biological replicates per condition derived from three independent experiments. Raw sequencing data were processed using the Galaxy Europe platform (usegalaxy.eu), including FASTQ extraction, read alignment, and downstream visualization. Paired-end reads were aligned to the GRCm39 mouse reference genome using HISAT2 with default parameters. Resulting alignments were filtered using BAM filtering criteria to retain only properly paired, mapped reads while excluding secondary alignments, low-quality reads, PCR duplicates, and reads overlapping blacklist regions (mm10-blacklist.v2.bed; ^44^). Differential gene expression analysis was conducted using the EdgeR package, applying a ≥ 1.5-fold change threshold and a false discovery rate (FDR)-adjusted *p*-value ≤ 0.05. Volcano plots were generated using the ggplot2 package in R. Gene clustering based on normalized counts was performed using the Morpheus matrix visualization and analysis tool. Functional enrichment analysis was carried out using ShinyGO (v0.85.1). Publicly available datasets were incorporated for comparative analyses ^26^ The ClusterProfiler package in R was used for the gene set enrichment analysis (GSEA).

### Statistics and Reproducibility

Flow cytometry data were analyzed using FlowJo v10.10.0 (BD Biosciences), and statistical analyses were performed with Prism v8 (GraphPad Software). Data are presented as mean ± SEM unless otherwise indicated. Comparisons between two groups were conducted using two-tailed paired or unpaired Student’s t tests, or the Mann–Whitney test when appropriate. For comparisons among three or more groups, one-way ANOVA was applied, while two-way ANOVA was used to analyze tumor growth curves. Multiple comparisons were corrected using Tukey’s or Sidak’s post hoc tests, as appropriate. A *p-value* < 0.05 was considered statistically significant.

## Data availability

Data associated with all figures and tables are provided in the Source data file and Supplementary files. The raw data generated in this study have been deposited in the Gene Expression Omnibus (GEO) database [https://www.ncbi.nlm.nih.gov/geo/] under the following accession number for RNA-seq data (https://www.ncbi.nlm.nih.gov/geo/query/acc.cgi?acc=GSE338454).

## Acknowledgments

J.D.Z.R. conducted this study to fulfill the requirements of the Programa de Doctorado en Ciencias Bioquímicas at Universidad Nacional Autónoma de México. We thank Dr. María Guadalupe García-Patiño for the critical reading and comments on the manuscript, Sandra Varela-Cardoso and Dr. Abraham Rosas Arellano for their technical support, and Jessica Lakshmi Prieto-Chávez at the Flow Cytometry core facility at Coordinación de Investigación en Salud in “Centro Médico Nacional, Siglo XXI”. We thank Dr. Laura Ongay Larios, MS. Minerva Mora Cabrera and Biol. María Guadalupe Códiz Huerta from the Molecular Biology Unit. We thank DVM. Claudia Rivera and DVM. Héctor Malagón from the Animal Facility. We thank the NIH Tetramer Core Facility (NIH Contract 75N93020D00005 and RRID: SCR_026557) for providing [64249 & 71638].

## Funding sources

This work was supported by the following grants: 303027 SECIHTI and PAPIIT IN220823 to P.L.L. J.D.Z.R., E.C.C., V.Y.G.R., D.B.R.R., and L.T.H. received a fellowship from SECIHTI 931997, 255287, 1228128, 935753, and 1101439, respectively. J.D.Z.R. received a fellowship from PAPIIT IN220823.

## Author contributions

P.L.L and J.D.Z.R. designed the project, analyzed the data, and prepared the manuscript. J.D.Z.R. conducted the experiments and analyzed the bioinformatic data. E.C.C., V.Y.G.R., C.M.M., D.B.R.R., J.L.R.B., J.O.C., A.I.S., C.G.E., F.R.S., and J.C.C.A. helped with the experiments. L.T.H. prepared the RNA-seq libraries. V.K provided the TIF1γ conditional-deficient mouse strain. R.F supported initial research support. P.L.L., J.D.Z.R., and E.C.C. discussed the data.

## Declaration of interests

The authors declare no competing interests.

## Notes

### Competing Interest Statement

The authors have declared no competing interest.

### Summary of Updates

Section authors have been updated to include an additional author: Mayra Furlan-Magaril

## References

1. W. Z. Mehal, S. Z. Sheikh, L. Gorelik, R. A. Flavell, TGF-β signaling regulates CD8+ T cell responses to high- and low-affinity TCR interactions. Int. Immunol. 17, 531–538 (2005).

2. S. Sanjabi, M. M. Mosaheb, R. A. Flavell, Opposing Effects of TGF-β and IL-15 Cytokines Control the Number of Short-Lived Effector CD8 + T Cells. Immunity 31, 131–144 (2009).

3. D. A. Thomas, J. Massagué, TGF-β directly targets cytotoxic T cell functions during tumor evasion of immune surveillance. Cancer Cell 8, 369–380 (2005).

4. A. P. Nath, et al., Comparative analysis reveals a role for TGF-β in shaping the residency-related transcriptional signature in tissue-resident memory CD8+ T cells. PLoS One 14, 1–16 (2019).

5. R. Ma, J. H. Sun, Y. Y. Wang, The role of transforming growth factor-β (TGF-β) in the formation of exhausted CD8+ T cells. Clin. Exp. Med. 24, 1–9 (2024).

6. S. S. Gabriel, et al., Transforming growth factor-β-regulated mTOR activity preserves cellular metabolism to maintain long-term T cell responses in chronic infection. Immunity 54, 1698–1714.e5 (2021).

7. Y. Hu, et al., TGF-β regulates the stem-like state of PD-1+ TCF-1+ virus-specific CD8 T cells during chronic infection. Journal of Experimental Medicine 219 (2022).

8. M. A. Travis, D. Sheppard, TGF-β activation and function in immunity. Annu. Rev. Immunol. 32, 51–82 (2014).

9. K. Chandiran, et al., SMAD4 and TGF-β are architects of inverse genetic programs during fate determination of antiviral CTLs. Elife 1–23 (2022). 10.7554/eLife.76457.

10. Y. Hu, Y.-T. Lee, S. M. Kaech, B. Garvy, L. S. Cauley, Smad4 Promotes Differentiation of Effector and Circulating Memory CD8 T Cells but Is Dispensable for Tissue-Resident Memory CD8 T Cells. The Journal of Immunology 194, 2407–2414 (2015).

11. R. Igalouzene, et al., SMAD4 TGF-β-independent function preconditions naive CD8+T cells to prevent severe chronic intestinal inflammation. J. Clin. Invest. 132, 1–19 (2022).

12. Y. Hu, Y.-T. Lee, S. M. Kaech, B. Garvy, L. S. Cauley, Smad4 Promotes Differentiation of Effector and Circulating Memory CD8 T Cells but Is Dispensable for Tissue-Resident Memory CD8 T Cells. The Journal of Immunology 194, 2407–2414 (2015).

13. J. Cao, et al., Smad4 represses the generation of memory-precursor effector T cells but is required for the differentiation of central memory T cells. Cell Death Dis. 6, 1–9 (2015).

14. X. Liu, et al., SMAD4, activated by the TCR-triggered MEK/ERK signaling pathway, critically regulates CD8+ T cell cytotoxic function. Sci. Adv. 8 (2022).

15. S. Dupont, et al., Germ-layer specification and control of cell growth by ectodermin, a Smad4 ubiquitin ligase. Cell 121, 87–99 (2005).

16. W. He, et al., Hematopoiesis Controlled by Distinct TIF1γ and Smad4 Branches of the TGFβ Pathway. Cell 125, 929–941 (2006).

17. J. M. Doisne, et al., iNKT cell development is orchestrated by different branches of TGF-β signaling. Journal of Experimental Medicine 206, 1365–1378 (2009).

18. S. Tanaka, et al., Trim33 mediates the proinflammatory function of Th17 cells. Journal of Experimental Medicine 215, 1853–1868 (2018).

19. E. Contreras-Castillo, et al., TIF1γ regulates stability of regulatory T cells during inflammation. Nat. Immunol. (2026). 10.1038/s41590-026-02603-6.

20. F. Ferri, et al., TRIM33 switches off Ifnb1 gene transcription during the late phase of macrophage activation. Nat. Commun. 6, 1–12 (2015).

21. H. Ali, et al., Cellular TRIM33 restrains HIV-1 infection by targeting viral integrase for proteasomal degradation. Nat. Commun. 10, 1–15 (2019).

22. X. Shen, et al., TRIM33 plays a critical role in regulating dendritic cell differentiation and homeostasis by modulating Irf8 and Bcl2l11 transcription. Cell. Mol. Immunol. (2024). 10.1038/s41423-024-01179-1.

23. I. Tiniakou, et al., Genome-wide screening identifies Trim33 as an essential regulator of dendritic cell differentiation. Sci. Immunol. 9, 1–19 (2024).

24. J. Xue, et al., Tumour suppressor TRIM33 targets nuclear β-catenin degradation. Nat. Commun. 6, 1–16 (2015).

25. M. E. Pipkin, et al., Interleukin-2 and Inflammation Induce Distinct Transcriptional Programs that Promote the Differentiation of Effector Cytolytic T Cells. Immunity 32, 79–90 (2010).

26. X. Meng, et al., FBXO38 mediates PD-1 ubiquitination and regulates anti-tumour immunity of T cells. Nature 564, 130–135 (2018).

27. S. Munir Alam, et al., Qualitative and Quantitative Differences in T Cell Receptor Binding of Agonist and Antagonist Ligands. Immunity 10, 227–237 (1999).

28. P. P. Yachi, J. Ampudia, T. Zal, N. R. J. Gascoigne, Altered Peptide Ligands Induce Delayed CD8-T Cell Receptor Interaction-a Role for CD8 in Distinguishing Antigen Quality. Immunity 25, 203–211 (2006).

29. M. O. Li, S. Sanjabi, R. A. Flavell, Transforming Growth Factor-β Controls Development, Homeostasis, and Tolerance of T Cells by Regulatory T Cell-Dependent and -Independent Mechanisms. Immunity 25, 455–471 (2006).

30. N. Zhang, M. J. Bevan, TGF-β signaling to T cells inhibits autoimmunity during lymphopenia-driven proliferation. Nat. Immunol. 13, 667–673 (2012).

31. E. Tu, et al., T Cell Receptor-Regulated TGF-β Type I Receptor Expression Determines T Cell Quiescence and Activation. Immunity 48, 745–759.e6 (2018).

32. V. Arumugam, et al., TCR signaling intensity controls CD8+ T cell responsiveness to TGF-. J. Leukoc. Biol. 98, 1–10 (2015).

33. M. Giroux, J.-S. Delisle, A. O’Brien, M.-J. Hébert, C. Perreault, T Cell Activation Leads to Protein Kinase Cθ-Dependent Inhibition of TGF-β Signaling. The Journal of Immunology 185, 1568–1576 (2010).

34. J. T. Tan, et al., IL-7 is critical for homeostatic proliferation and survival of naïve T cells. Proceedings of the National Academy of the United States of America 98, 8732–8737 (2001).

35. M. Y. Kimura, et al., IL-7 signaling must be intermittent, not continuous, during CD8 + T cell homeostasis to promote cell survival instead of cell death. Nat. Immunol. 14, 143–151 (2013).

36. M. O. Li, S. Sanjabi, R. A. A. Flavell, Transforming Growth Factor-β Controls Development, Homeostasis, and Tolerance of T Cells by Regulatory T Cell-Dependent and -Independent Mechanisms. Immunity 25, 455–471 (2006).

37. T. C. Wirth, J. T. Harty, V. P. Badovinac, Modulating numbers and phenotype of CD8+ T cells in secondary immune responses. Eur. J. Immunol. 40, 1916–1926 (2010).

38. M. R. Jenkins, A. Tsun, J. C. Stinchcombe, G. M. Griffiths, The Strength of T Cell Receptor Signal Controls the Polarization of Cytotoxic Machinery to the Immunological Synapse. Immunity 31, 621–631 (2009).

39. S. Yao, et al., Interferon regulatory factor 4 sustains CD8+ T cell expansion and effector differentiation. Immunity 39, 833–845 (2013).

40. K. Wei, et al., TRIM28 is an essential regulator of three-dimensional chromatin state underpinning CD8+ T cell activation. Nat. Commun. 16, 1–20 (2025).

41. A. M. Intlekofer, et al., Effector and memory CD8+ T cell fate coupled by T-bet and eomesodermin. Nat. Immunol. 6, 1236–1244 (2005).

42. Z. Liu, et al., Cutting Edge: Transcription Factor BCL6 Is Required for the Generation, but Not Maintenance, of Memory CD8+ T Cells in Acute Viral Infection. The Journal of Immunology 203, 323–327 (2019).

43. Q. Sun, et al., BCL6 promotes a stem-like CD8 + T cell program in cancer via antagonizing BLIMP1. Sci. Immunol. 8, 1–17 (2023).

44. H. M. Amemiya, A. Kundaje, A. P. Boyle, The ENCODE Blacklist: Identification of Problematic Regions of the Genome. Sci. Rep. 9, 1–5 (2019).

